# Coalescing nephron and ureteric bud progenitors potentiates nephrogenesis in recellularized kidney scaffolds

**DOI:** 10.64898/2026.06.24.733560

**Authors:** Ashwani Kumar Gupta, Ekta Minocha, Jiao-Jing Wang, Zhenxiao Tu, Zheng J Zhang, Jason A. Wertheim

## Abstract

Bioengineered, transplantable kidney tissue using decellularized scaffolds offers a promising strategy to overcome the shortage of donor kidneys that limits organ transplantation for patients with end stage renal disease. These kidney scaffolds retain essential extracellular matrix architecture, providing a biologically active niche for recellularization. Successful generation of bioengineered kidney tissues includes enhanced patent vasculature and mature, functional nephrons with collecting ducts. Here, we report the development of engineered kidney tissue consisting of reconstituted kidney scaffolds and human pluripotent stem cell-derived nephron and ureteric bud progenitors. Structural analysis of recellularized kidney scaffolds showed advanced nephron structures that became more mature and exhibited interconnected nephron and collecting ducts. *In vivo* engraftment of reconstituted kidney scaffolds in mice led to vascularization, maturation, and secretory function. Notably, mouse-graft vascular anastomosis was evident with erythrocytes present in vasculature and nephron-secreted proteins detected in mouse urine, indicating functional integration. This approach demonstrates the feasibility to generate advanced bioengineered kidney tissues that offer a versatile platform for disease modeling, drug screening, and regenerative medicine.

**Graphical abstract:** 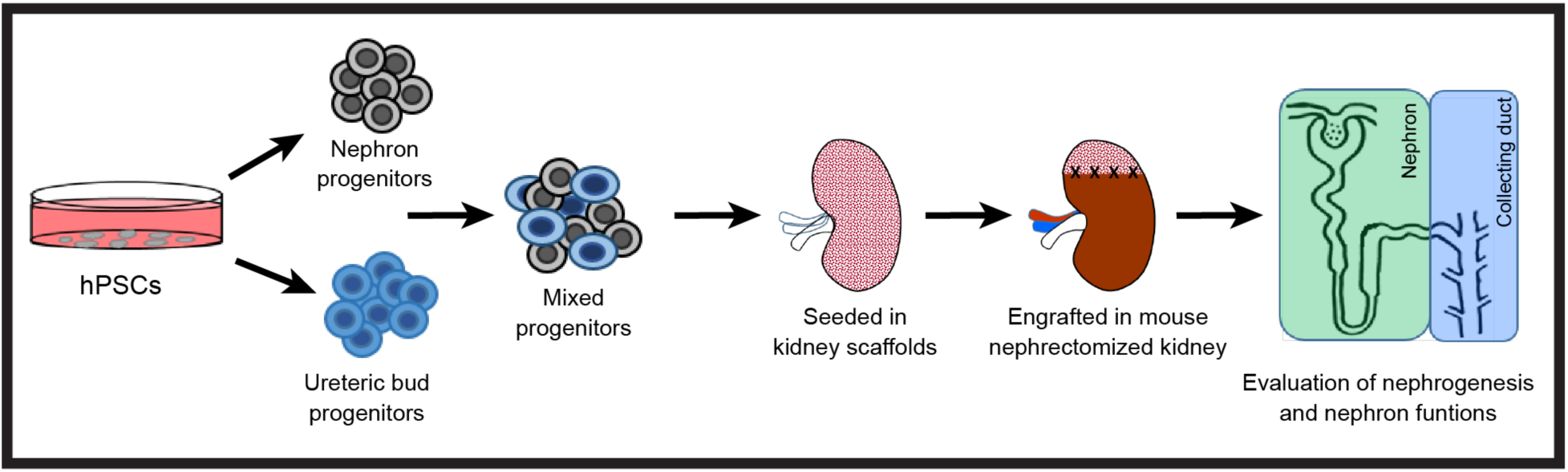

## Introduction

End Stage Renal Disease (ESRD) is a global healthcare concern, affecting ∼10% of the worldwide population^1^. The only therapeutic options for patients with ESRD are dialysis and kidney transplantation, although the limitations of these therapies impact their long-term success^2^. In recent years, kidney tissue engineering has emerged with a potential to repair or replace damaged kidney tissues^3,4^. The ability to generate kidney tissues has improved, with directed differentiation of human pluripotent stem cells (hPSCs) into nephron progenitors (NPs) and ureteric bud progenitors (UBPs) that further mature to form nephron-like epithelial structures^5–8^.

During organogenesis, mesenchyme to epithelial transition leads to nephron epithelialization, induced by reciprocal interaction between the ureteric bud (UB) derived from anterior intermediate mesoderm (AIM) and the metanephric mesenchyme (MM) derived from posterior intermediate mesoderm (PIM)^9^. MM induces ureteric buds to undergo branching morphogenesis and give rise to collecting ducts^10,11^. Stromal progenitors surround the nascent nephron and produce extracellular matrix (ECM), which provides 3D structure and biochemical cues critical for promoting and maintaining cell differentiation^12,13^.

Native organ scaffolds maintain the organ specific intrinsic architecture, ECM proteins, vascular structures, and associated signaling factors. These features orchestrate essential cellular behaviors like adhesion, migration, differentiation, and gene regulation in the matrix. Consequently, native organ-derived scaffolds offer both the structural framework and instructive environment necessary for effective organ regeneration^14^. Given the central role of the ECM, decellularization of whole organs have been utilized to generate native organ scaffolds that may be reconstituted with cells of renal lineages^15,16^. Our group previously reported a decellularization method for rat kidneys that maintains native microarchitecture and supports kidney bioengineering applications^17^. While several studies have attempted to generate vasculature and renal tissues using kidney scaffolds, progress remains limited. Tajima et al, 2022 reported nephron induction and regeneration without fibrillogenesis when a porcine kidney-derived extracellular matrix was sutured onto the surface of partial nephrectomized kidney^14^. Another study by Song et al, 2013 demonstrated limited renal tissue formation and function following recellularization of rat kidney scaffolds with rat neonatal kidney cells and human umbilical venous endothelial cells (HUVEC)^18^. Additional studies have also shown encouraging but incomplete outcomes in vascularization, renal tissue formation, and functional recovery^4,19–22^. Despite these advances, recapitulation of *in vivo*-like renal tissue architecture and function has not been fully achieved, and no study has reconstituted kidney scaffolds with combinations of NPs and UBPs derived from hPSCs and assessed function *in vivo*.

Here, we generated NPs and UBPs by differentiating hPSCs separately and subsequently combined these progenitors to develop kidney organoids. We demonstrate that combining these progenitors enhances tubular density and promotes more robust nephron formation within organoids. We further evaluated this approach to recellularize native acellular kidney scaffolds by seeding them with mixed progenitors and implanting the recellularized kidney scaffolds *in vivo*. This study investigates how coalescing nephron and ureteric bud progenitors influence nephrogenesis, nephron secretory function and vascularization in native kidney scaffolds and after engraftment of these bioengineered constructs into partially nephrectomized animal model.

## Results

### Generation and characterization of nephron and ureteric bud progenitors

During kidney organogenesis, the crosstalk between nephron and ureteric bud progenitors within the ECM microenvironment leads to formation of nephron and collecting ducts. We therefore hypothesized that combining NPs and UBPs in acellular native kidney scaffolds may potentiate a higher order of nephrogenesis. We followed the Morizane et al. 2015 and Tsujimoto et al. 2020 protocols to differentiate hPSCs into NPs and UBPs, respectively^23,24^, and combined them to construct a nephrogenic-like microenvironment. To provide a synchronous timeframe for reference (day 0, when UBP differentiation started), the differentiation of NPs began three days prior (day -3) to the differentiation of hPSCs into UBPs (Fig. 1a). During NPs differentiation, hPSCs lost the pluripotent gene OCT4 and NANOG expression on day 1 and robustly expressed genes of the mesoderm, T and TBX6. On day 4, the expression of mesoderm genes decreased as expected and cells differentiated into OSR1^+^ HOXD11^+^ PIM (Fig. 1b,c). On day 6, PIM cells differentiated into SIX2, SALL1, WT1 and GDNF-expressing NPs (Fig. 1b,c). On day 1 of UBPs differentiation, hPSCs lost pluripotent gene OCT4 and NANOG expression and began to express mesodermal genes, T and TBX6. On day 4, mesodermal genes were downregulated and cells differentiated into OSR1^+^ LHX1^+^ AIM (Fig. 1d,e). On day 6, AIM cells were differentiated into GATA3, PAX2, ETV5 and WNT9B-expressing UBPs (Fig. 1d,e). The expression of early nephron and UB specific progenitor markers indicate that hPSCs differentiated into NPs and UBPs.

**Fig 1.**
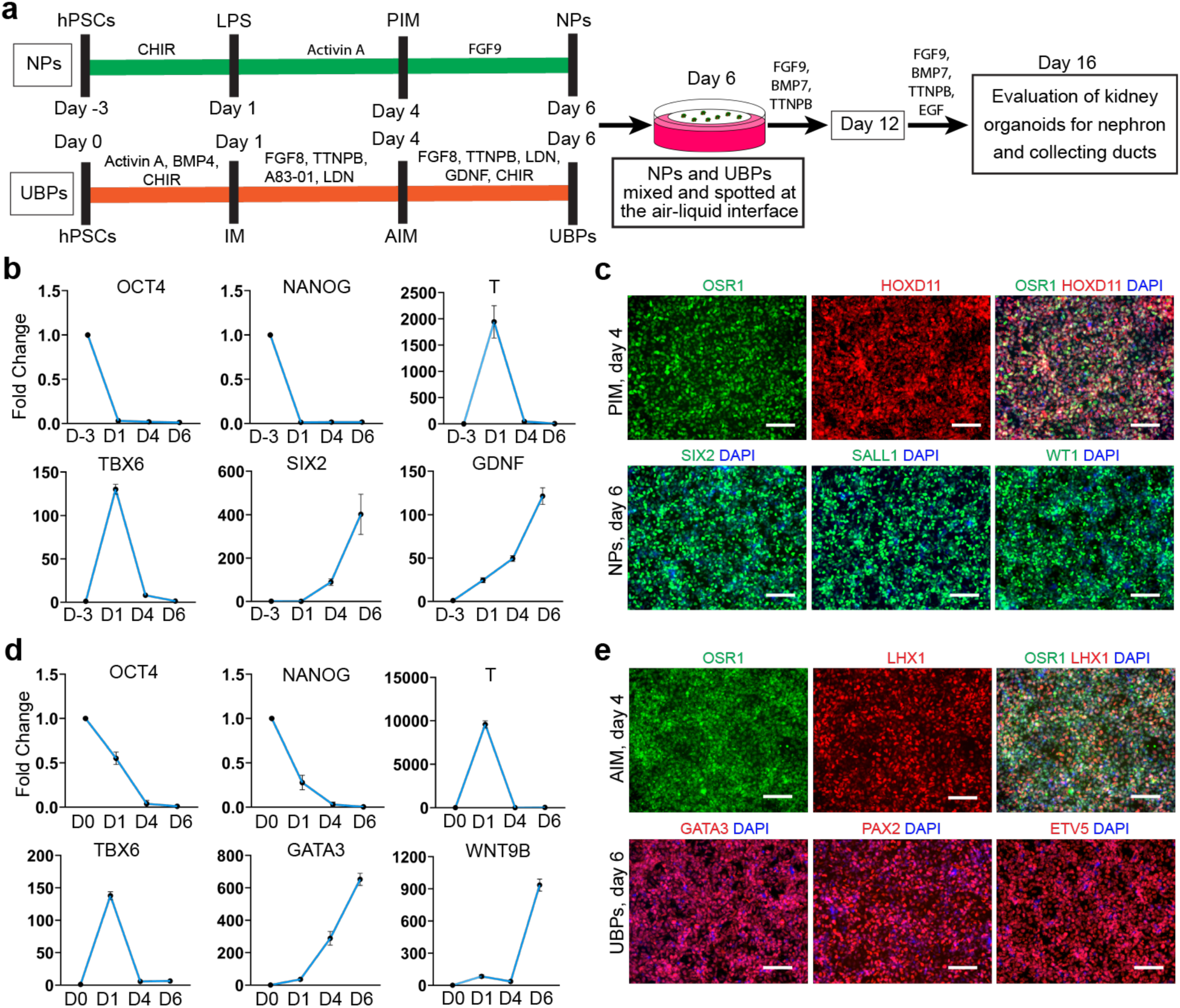
Schematic illustration showing generation and characterization of nephron and ureteric bud progenitors. **a.** Schematic diagram shows the stepwise differentiation strategy of hPSCs into NPs and UBPs, and generation of kidney organoids. **b.** Changes in gene expression of pluripotency, mesoderm and NPs-related makers during NPs differentiation. (n = 3 independent biological replicates per time-point). **c.** Representative immunofluorescence images shows the differentiation of hPSCs into PIM on day 4 and differentiation into NPs expressing SIX2, SALL1 and WT1 on day 6. Scale bars 100µm. **d.** Changes in gene expression of pluripotency, mesoderm and UBPs related makers during UB differentiation. (n = 3 independent biological replicates per time-point). **e.** Representative immunofluorescence images shows differentiation of hPSCs into AIM and differentiation into UBPs expressing GATA3, PAX2 and ETV5. Scale bars 100µm.

### Combining nephron and ureteric bud progenitors leads to enhanced nephron formation in kidney organoids

Kidney organoids were generated by mixing NPs and UBPs at a ratio 3:1 and aggregated at the air-liquid interface. During kidney development, NPs and UBPs exist together where UB secrete WNT9B that induce WNT-dependent epithelialization of NPs. To activate WNT signaling, GSK3β inhibitor CHIR99021 is used widely to epithelialize NPs in kidney organoid protocols^23,25^. In our protocol, the effect of combining UBPs with NPs omitted the need for CHIR99021, which is believed to have multiple effects other than inducing WNT-dependent epithelialization^26^. The combination of both progenitors resulted in kidney organoids with tightly packed epithelial structures compared to organoids generated from NPs only (Fig. 2a). Kidney organoids generated from mixing of NPs and UBPs had significantly high area fraction of expressed molecular markers NPHS1, LTL, CDH1, KRT8 and PECAM1 in addition to significantly upregulated gene expression indicative of discrete nephron segments and collecting ducts (Fig. 2b and Supplementary Fig. S2). Further, immunofluorescence staining revealed that kidney organoids generated from mixed progenitors were packed with WT1-expressing podocytes clusters, LTL positive proximal tubules, PECAM1-expressing endothelial networks, POU3F3-expressing distal tubules, and CALB1, PAX2, HOXB7, RET, KRT8-expressing collecting ducts (Fig. 2c). Overall, mixing nephron progenitors with UB progenitors resulted in robust epithelialization, presence of collecting ducts, and rich nephrogenesis within the kidney organoids, without the need for WNT stimulators.

**Fig 2.**
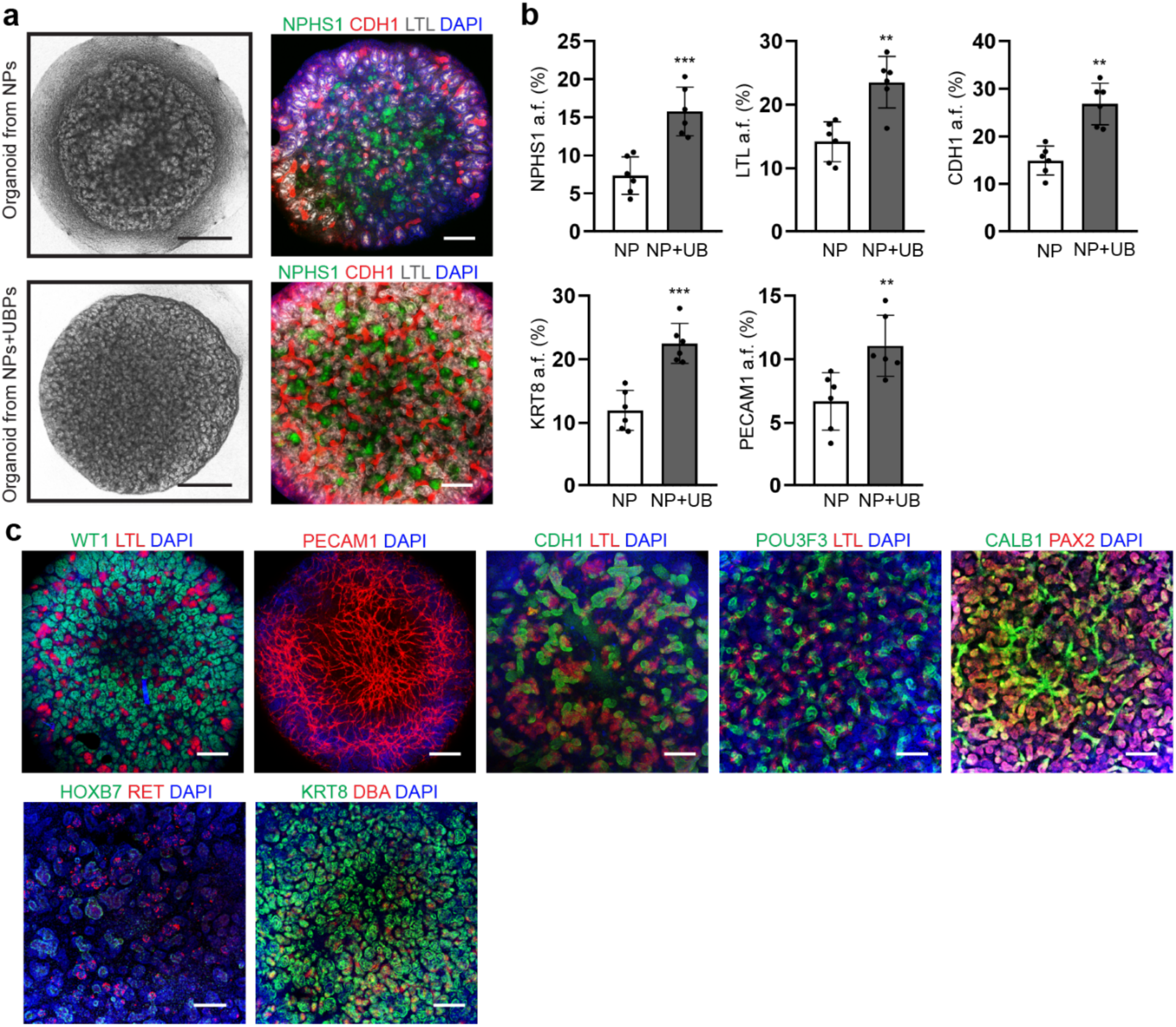
Enhanced nephrogenesis in kidney organoids after combination of nephron and ureteric bud progenitor populations. **a.** Representative stereo microscope (scale bars 500µm) and immunofluorescence images (scale bars 200µm) of kidney organoids generated from NPs only, and a combination of NPs and UBPs. **b.** Image J quantification of stained area fraction for NPHS1, LTL, CDH1, KRT8 and PECAM1. (n = 6 quantified fields from three independent biological replicates). Data presented as mean ± s.d. \*\**P* < 0.01; \*\*\**P* < 0.001. **c.** Representative immunofluorescence images shows that combining NPs and UBPs (at a ratio 3:1) leads to kidney organoids filled with podocytes, proximal and distal tubules, endothelial networks, and collecting ducts. Scale bars 200µm.

To provide greater flexibility, we tested whether both progenitor populations preserve their phenotype and potential for nephrogenesis after cryopreservation. On day 6 of differentiation, both progenitor populations were cryopreserved and stored in liquid nitrogen. We successfully revived both progenitor populations that had been cryopreserved for more than 12 months. Revived cells were cultured for 24h, and immunofluorescence staining showed that NPs expressed SIX2 and SALL1, while UBPs expressed GATA3 and PAX2. Further, kidney organoids generated from the combination of both thawed progenitor populations contained tightly packed tubular clusters that were filled with NPHS1^+^ KRT8^+^ and LTL^+^ structures. This confirms that NPs and UBPs retain their progenitor state and differentiation potential even after cryopreservation (Supplementary Fig. 1a-f).

### Nephron and UB progenitors promoted vascularization with functional nephrogenesis in recellularized kidney scaffolds

Once we confirmed that combining nephron and ureteric bud progenitors potentiates nephrogenesis in organoids, we sought to use this strategy to recellularize acellular kidney scaffolds. Rat native kidneys were cannulated, harvested and decellularized (Fig. 3a,b and Supplementary Fig. S3) by following our previously reported protocol^27^. This resulted in removal of the cellular component of the kidney, which was confirmed by removal of ∼95% of DNA within the kidney. We found that acellular scaffolds contained 75.49±18.93 ng DNA/mg of wet tissue whereas native kidneys contained 1833±173.2 ng DNA/mg of wet tissue (Fig. 3c). Further, native kidney scaffolds retain considerable amount of extracellular matrix (ECM) proteins, such as fibronectin (9.83±1.77 ng/25mg of tissue) and GAGs (228.5±50.28 µg/25mg of tissue) (Fig. 3d,e).

**Fig 3.**
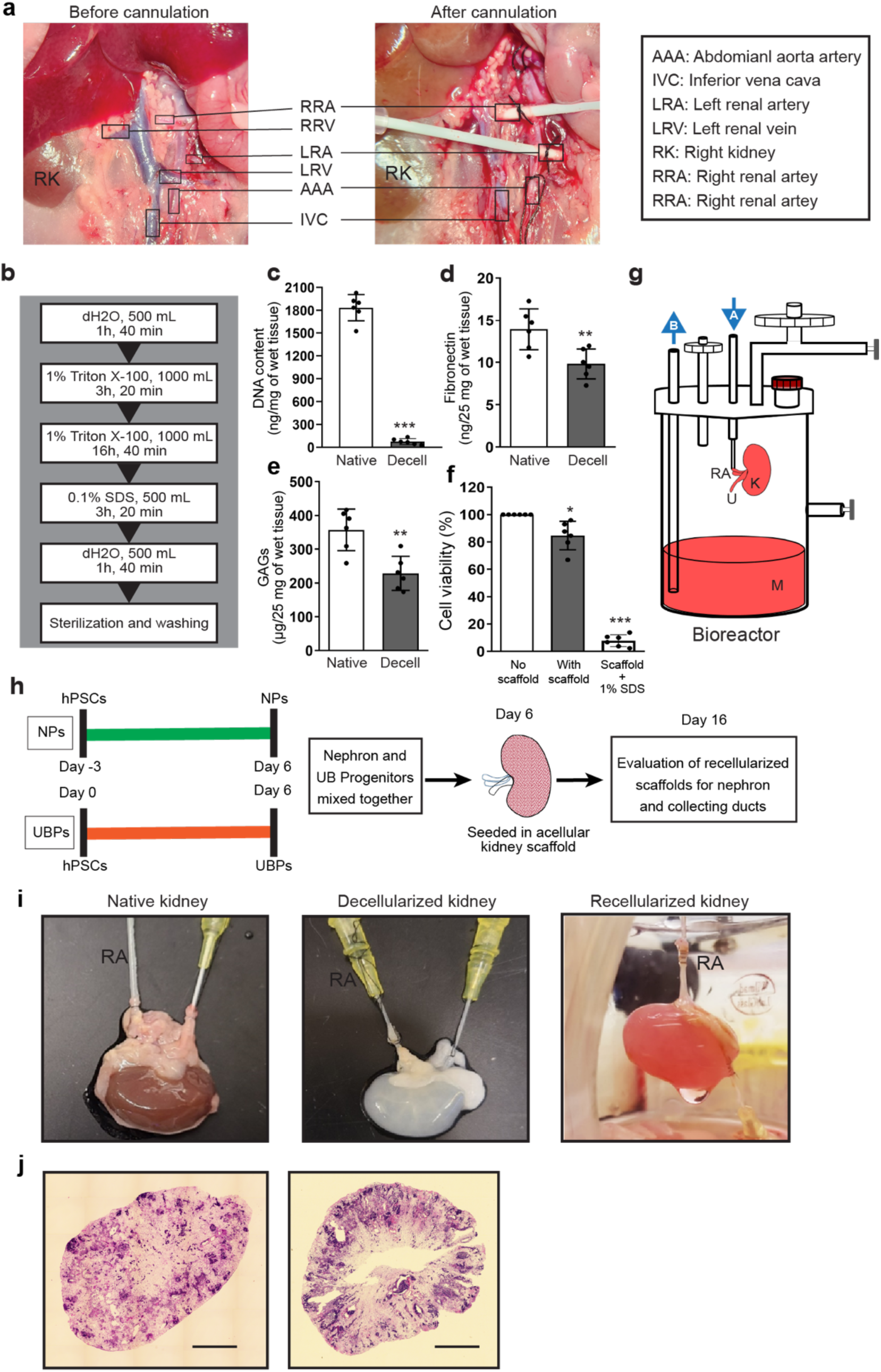
Acellular rodent native kidney scaffold preparation and recellularization with a combination of nephron and ureteric bud progenitors. **a.** Images show renal artery cannulation of the rat kidney. **b.** Schematic diagram shows stepwise decellularization of rat native kidney. **c.** Quantification of DNA content and ECM components including **d.** fibronectin, **e.** GAGs in native and decellularized kidney scaffolds. **f.** Quantification of cell viability (%) after seeding NPs and UBPs (at a ratio 3:1) in culture wells (No scaffold), in scaffold (with scaffold) and in scaffold with recellularization media containing 1% SDS as a negative control. (n = 6 cultures from three independent biological replicates per group). Data presented as mean ± s.d. \*\**P* < 0.01, ***P < 0.001. **g.** Schematic diagram shows recellularization bioreactor assembly with media (M) inlet (A), media outlet (B), kidney (K), renal artery (RA) and ureter (U). **h.** Schematic diagram shows the recellularization strategy with a combination of hPSCs after differentiation into NPs and UBPs (at a ratio 3:1). **i.** Images depict the rat kidney before and after decellularization, and after recellularization. **j.** Hematoxylin and Eosin staining shows sagittal and transverse sections of recellularized kidney scaffolds. Scale bars 2mm.

Sterilized kidney scaffolds were seeded with mixed NPs and UBPs at a ratio of 3:1. MTT assay was performed to evaluate the cell viability of seeded cells after 24 hours of culture in the bioreactor. We observed that >80% of cells were viable after growth in the bioreactor, whereas <10% cells were viable in a negative control containing 1% SDS in recellularization media, in comparison to progenitors cultured in the absence of a scaffold (Fig. 3f). After 10 days of culture within the bioreactor, kidney scaffolds were evaluated for nephron, collecting ducts and vascularization (Fig. 3g-i and Supplementary Fig. S4). Immunofluorescence staining revealed SIX2-expressing nephron progenitors surrounding KRT8 and DBA-expressing cells of UB lineage (Supplementary Fig. S5a), apparently establishing nephrogenic zone-like structures that have been reported to give rise to epithelial nephron structures because of reciprocal crosstalk between nephron and ureteric bud progenitors^25^. The interaction between nephron and UB progenitors resulted in nephron patterning with collecting duct networks throughout the scaffold. NPHS1-expressing glomerular structures were found connected to LTL positive proximal tubules. PECAM1-expressing endothelial networks invaded PODXL and WT1-expressing glomeruli. Renin-expressing juxtaglomerular (JG) cells were present adjacent to PODXL-expressing glomeruli. Proximal tubules in the scaffold expressed NaKATPase, UMOD expression was detected in tubules suggesting loop of Henle, and distal tubules expressed CDH1 and POU3F3. Furthermore, collecting ducts expressed definitive markers such as KRT8, GATA3 and DBA with connection to UPKII-expressing tubules (Fig. 4a). To assess if endothelial networks have patent lumens, reconstituted scaffolds were perfused with fluorescent Ulex Europaeus Agglutinin I (UEAI). The luminal detection of UEA1 in PECAM1-expressing endothelial networks confirmed vessel maturation, emphasizing the benefits of flow within the bioreactor (Fig. 4b). During blood pressure and fluid balance regulation, arginine vasopressin (AVP) indirectly stimulates AQP2 transcription and aldosterone release from the adrenal gland, which diffuses into principal cells and triggers rapid translocation of AQP2 from the basal to apical membrane of collecting ducts^28,29^. To test collecting duct maturity and AQP2 translocation ability, we treated recellularized scaffolds with 10 nM Aldo and 10 nM AVP for 48 hours in the perfusion bioreactor. The treatment with aldosterone and vasopressin resulted in AQP2 translocation from the basal to apical membrane of KRT8-expressing collecting ducts. (Fig. 4c). Further, 24h bioreactor perfused recellularization media was evaluated for the secretion of renin and UMOD by ELISA, which showed significantly higher levels of secreted renin (3494±191 pg ml^-1^) and UMOD (1526±109 pg ml^-1^) compared to recellularization media used to perfuse acellular scaffolds (Fig. 4d). Next, to evaluate function by conversion of 25-hydroxyvitamin D3 into its active form 1, 25-dihydroxyvitamin D3 via 1-α-hydroxylase in the proximal tubule^30^, the recellularized kidney scaffold was treated with 25-hydroxyvitamin D3 for 24h. A significantly higher level of 1, 25-dihydroxyvitamin D3 (160±8 pg mL^-1^) was detected by ELISA in the treated media compared to the untreated control (Fig. 4d). These results suggest that the recellularization of kidney scaffolds with NPs and UBPs resulted in more mature endothelial network and functional JG cells, proximal tubule, loop of Henle and collecting ducts. In summary, these tests confirm that the recellularized scaffolds are comprised of nephron segments and collecting ducts that have functional attributes.

**Fig 4.**
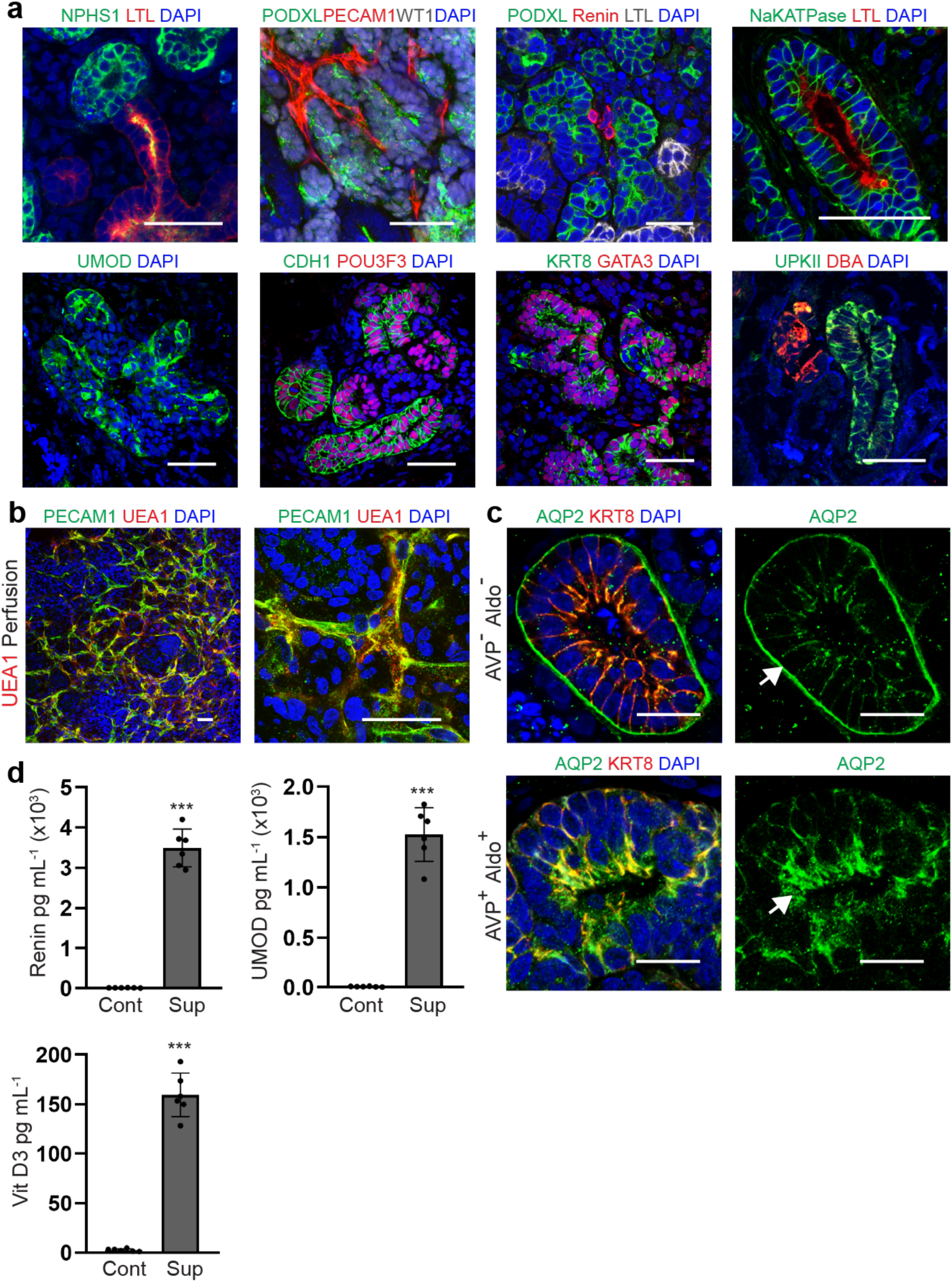
Nephrogenesis in recellularized kidney scaffolds and secretory function of nephrons. **a.** Representative immunofluorescence images of recellularized scaffolds shows nephron and collecting duct structures with segment specific expression of different markers including glomerulus and proximal tubule interconnection, and PECAM1^+^ vascular invasion in PODXL^+^ glomerular structures. Scale bars 50µm. **b.** Representative immunofluorescence images of recellularized scaffolds shows co-localization and luminal detection of perfused DyLight594-conjugated UEA1 in PECAM1^+^ endothelial network. Scale bars 50µm**. c.** Representative immunofluorescence images shows translocation of AQP2 from basal to apical site of KRT8-expressing collecting duct after perfusion with vasopressin (AVP) and aldosterone (Aldo) for 48h in recellularized scaffolds. Scale bars 20µm **d.** Graphs depict quantification of secretory proteins in the bioreactor perfusion media by ELISA. Recellularization resulted in secretion of renin and uromodulin (UMOD) during 24h of perfusion with recellularization media (supernatant noted as Sup) in comparison to acellular scaffolds perfused with recellularization media as a control (Cont). Further, 1, 25-dihydroxyvitamin D3 (Vit D3) was detected in the recellularized scaffold perfused with recellularization media (Sup) for 24h and treated with 25-hydroxyvitamin D3 in comparison to acellular scaffolds perfused with recellularization media as a control (Cont). n = 6 recellularized scaffolds from three independent biological replicates. Data presented as mean ± s.d. \*\*\**P* < 0.001.

### *In vivo* engraftment promotes vascularization, maturation and secretory function of nephrons

We sought to investigate the vascularization, cellular maturation and secretory function of reconstituted kidney scaffolds *in vivo*. Maturity and host-derived vascularization of engrafted kidney tissues have been reported, but fusion of engrafted epithelial tubules has not yet been well demonstrated. Epithelial fusion is necessary for urine outflow from the graft. To improve structural organization at a cellular level and provide a structure for implantation, we recellularized acellular kidney scaffolds that maintain the ECM architecture as a template^27^. We added NPs and UBPs to the scaffold that were incubated within a perfused bioreactor as before, now to investigate the presence of interconnected nephron and collecting ducts after engraftment. Partial nephrectomy is a surgical procedure used for different disease conditions (such as removal of small kidney tumors), and we used it here to investigate stimulation of matrix-assisted self-repair of remnant renal tissue, which could further assist maturation of reconstituted kidney scaffolds^14^. To provide this interactive tissue environment we engrafted recellularized kidney scaffolds onto the partially nephrectomized remnant kidney of NSG mice (Fig. 5a). We found that the size of the graft increased over three weeks of engraftment (Fig. 5d). Vascularization of tissue is important for the survival and maturation of grafted tissue. We observed visible blood vessels within the graft at the time of graft recovery (Fig. 5e).

**Fig 5.**
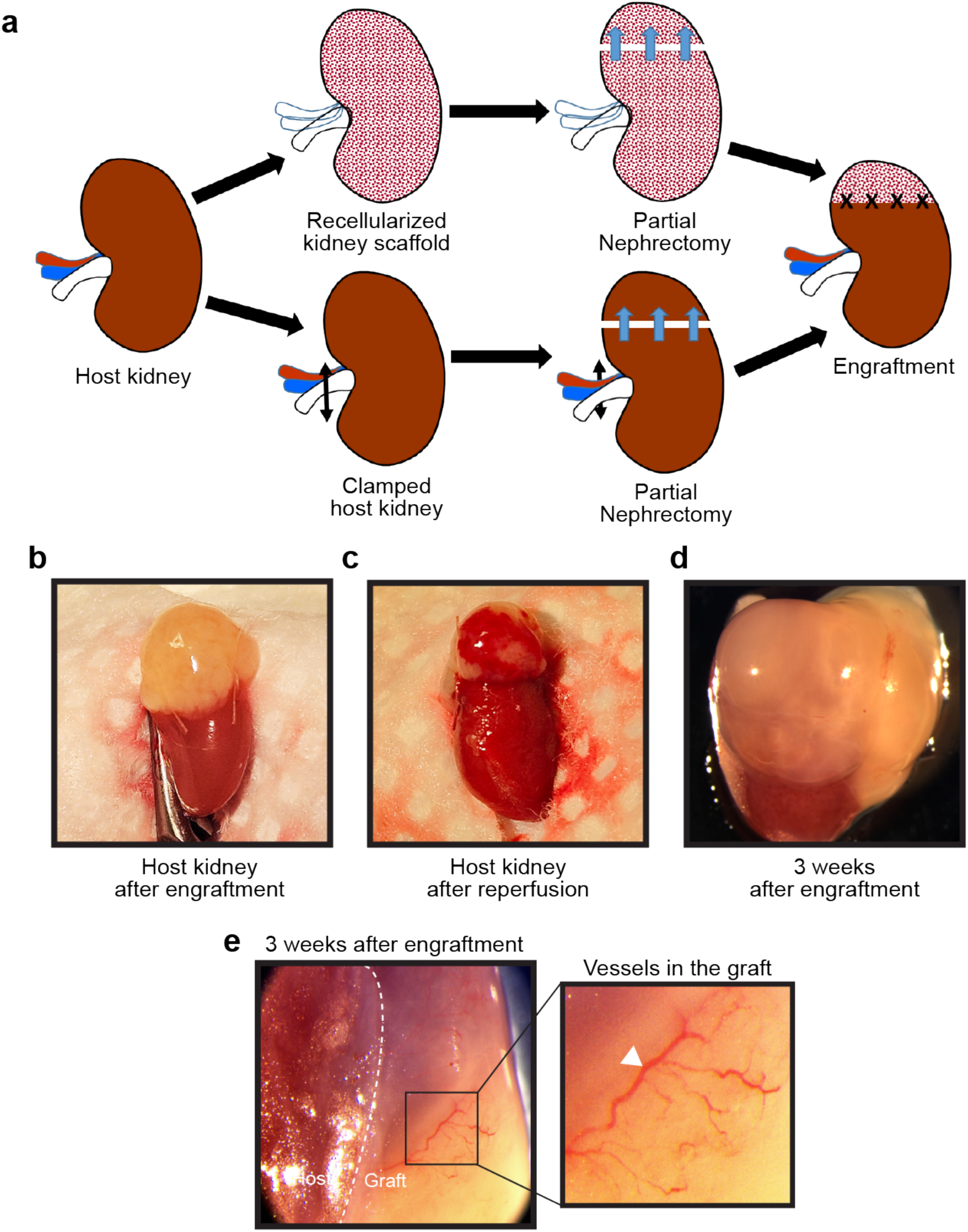
Schematic illustration showing engraftment of recellularized scaffold in partially nephrectomized mice kidney and appearance of graft after three weeks of engraftment. **a.** Schematic diagram shows engraftment of the recellularized kidney scaffold to the cut surface of the remnant kidney of NSG mice after partial nephrectomy. **b.** Image shows engrafted host kidney after scaffold implantation, **c.** after blood reperfusion and **d.** three weeks after engraftment. **e.** Image shows visible vessels after three weeks of engraftment.

To assess the origin of endothelial networks, we used mouse and human specific PECAM1 antibodies. Immunofluorescence staining of engrafted kidney vibratome sections revealed extensive vascularization (Fig. 6a) and host-graft vascular anastomosis in the graft. Further, staining for markers of red blood cells (RBCs) revealed the presence of RBCs in both human- and mouse-specific vessels and glomerular structures. Invasion of PECAM1-expressing endothelial vessels into the presumptive glomeruli forming capillary tuft like structures, were also observed (Fig. 6b). Further, immunofluorescence staining revealed CDH1^+^ epithelialization (Fig. 6c) and structural assessment revealed that the graft was filled with complex nephron structures including podocytes clusters expressing NPHS1, NPHS2, and PODXL. The podocyte clusters or presumptive glomerular structures showed connection with LTL positive proximal tubules. PDGFRβ-expressing mesangial cells were also found within PODXL-expressing glomerular structures. The basolateral expression of NaKATPase and apical staining for LTL suggest that proximal tubules were polarized within the graft and connected to NKCC2 or UMOD-expressing loop of Henle. Additionally, luminal expression of LRP2 was detected in LTL positive proximal tubules, and juxtaglomerular (JG) cells were detected at the base of PDXL-expressing glomeruli, suggesting development of JG apparatus-like structures (Fig. 6d). Next NCC-expressing distal tubule were found connected with DBA positive branched collecting ducts (Supplementary Fig. S5b,c). The branched collecting ducts also showed expression of principal cell markers AQP2, SCNN1A, SCNN1B, and SCNN1G. Further, microtubules expressing Ac-α-tubulin were found in KRT8 and DBA-positive collecting ducts (Fig. 6d). To determine origin of cellular components in the graft derived from human or mouse, we stained tissue sections for human nuclear antigen (HuNu). Staining with HuNu antibody confirmed that nephron structures including glomerulus, proximal tubules, distal tubules and collecting ducts in the graft were derived from human engrafted tissues along with PDGFRβ expressing stromal cells around tubules (Fig. 7a,b).

**Fig 6.**
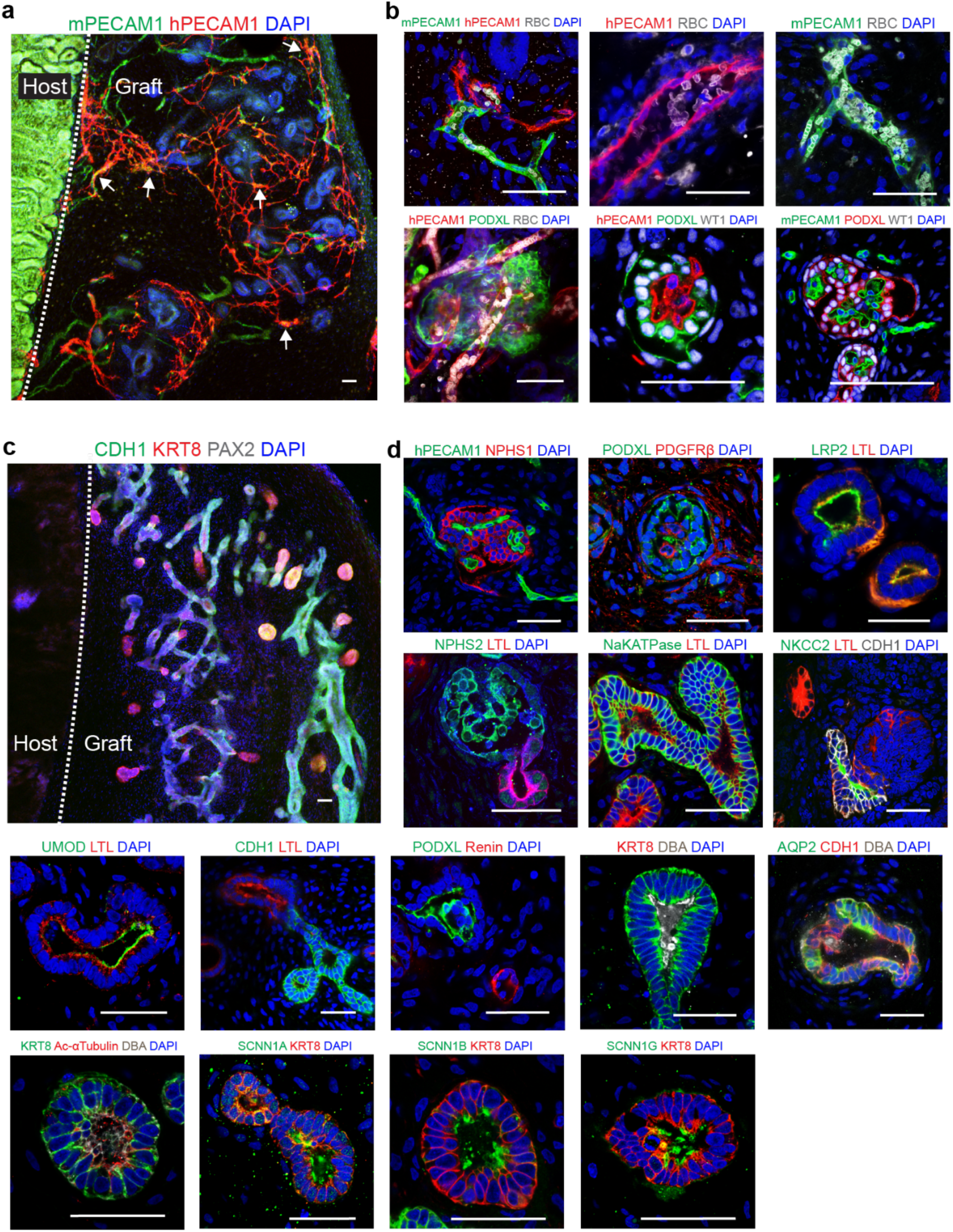
Reconstituted grafts lead to vascularization and enhanced nephron formation after engraftment. **a.** Representative immunofluorescence image depicts extensive vascularization, invasion of mouse (mPECAM1^+^) vessel into the graft and anastomosis with human (hPECAM1^+^) vessels (white arrow). **b.** Representative high magnification immunofluorescence images show host and graft vascular anastomosis and the presence of RBCs in the graft. Mouse RBCs were detected in both mouse and human-specific vessels and formation of glomerular capillary tuft like structures in PODXL^+^ and WT1^+^ human presumptive glomeruli in the graft. **c.** Representative immunofluorescence images show CDH1^+^ epithelialization in the graft. **d.** Invasion of hPECAM1^+^ vascular networks were identified in NPHS1^+^ glomerular structures. Mesangial cells expressing PDGFRβ were also detected in the glomerulus. Proximal tubules became more mature after engraftment and showed luminal expression of LRP2 and Na^+^K^+^ATPase, with connections to NPHS2^+^ glomeruli, CDH1^+^ NKCC2^+^ and UMOD^+^ loop of Henle (LOH) along with Renin^+^ Juxtaglomerular (JG) cells. Representative immunofluorescence images show mature collecting ducts expressing KRT8, AQP2, SCNN1A, SCNN1B, SCNN1G and presence of Ac-αTubulin^+^ luminal microvilli. Scale bars 50µm.

**Fig 7.**
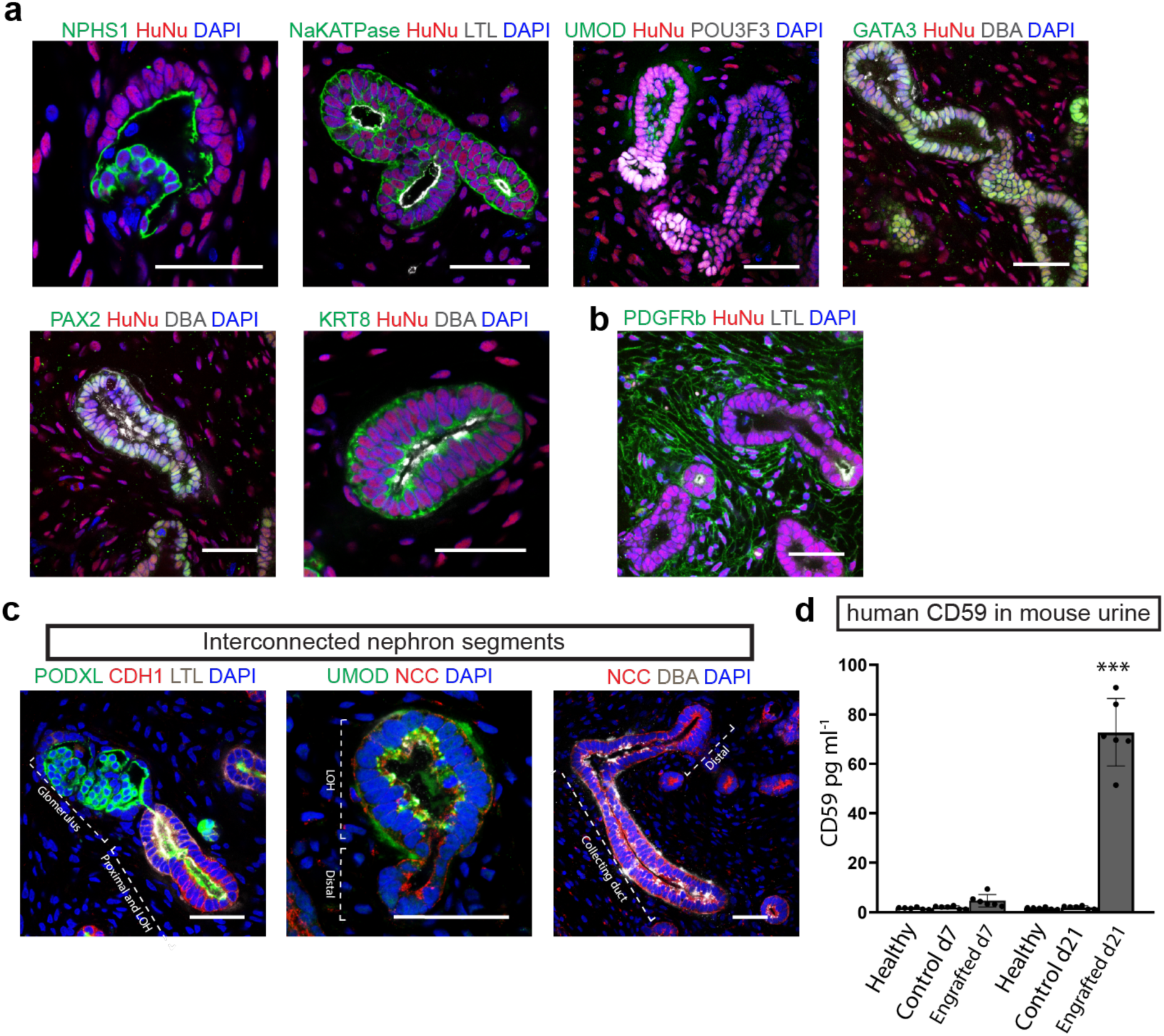
Graft characterization for human specific nephron structures and integrated nephron segments. **a.** To characterize nephron structures, human specific antibody HuNu was used. Representative immunofluorescence stains reveal that the cells of NPHS1^+^ glomerular structures, NaKATPase^+^ LTL^+^ proximal tubules, UMOD^+^ POU3F3^+^ loop of Henle (LOH), GATA3^+^ DBA^+^ PAX2^+^ KRT8^+^ collecting ducts are positive for HuNu, which indicates that these structures within the graft are derived from the human cells of the recellularized scaffold. **b.** PDGFRβ^+^ HuNu^+^ stromal cells were detected around tubules in the graft suggest that the stromal cells were derived from human graft. **c.** Further, representative immunofluorescence images show interconnected nephron segments: PODXL^+^ glomerulus, LTL^+^ proximal tubule, UMOD^+^ LOH, NCC^+^ distal tubule and DBA^+^ collecting duct. Scale bars 50µm. **k.** Human protein CD59 was detected at a significantly higher level in the urine of engrafted mice on day (d) 21 (engraftment) in comparison to mice implanted with an acellular scaffold (control) or healthy mice urine (healthy), (n = 6 mouse per group from three independent biological replicates). Data presented as mean ± s.d. \*\**P* < 0.01.

The kidneys play a vital role in maintaining total body water homeostasis, due to its filtration and secretory functions that require full length of nephrons connected to collecting ducts. Immunofluorescence staining of tissue sections after three weeks of engraftment revealed that presumptive glomerular structures connected to LTL positive proximal tubules and further linked to CHD1-expressing loop of Henle. UMOD-expressing LOH was found connected to NCC-expressing distal tubules, which was further connected to DBA positive collecting ducts, altogether indicating that nephrons became mature and contained interconnected segments within the graft (Fig. 7c). In addition to histological evaluation, we further validated nephron epithelial tubule fusion by detection of graft-derived proteins: CD59, renin and UMOD in mouse urine. CD59 is expressed in the glomerulus and proximal tubule reported to protect from complement-mediated injury, renin is a component of renin-angiotensin system (RAS) and is secreted by JG cells and UMOD is secreted from distal nephron and is an anti-bacterial agent in urine^31^. Recently, Chuang et al., reported a method for identification of human urinary biomarkers (CD59 and UMOD) in mouse urine, which can indicate tubule fusion and function of proximal and distal nephrons^31^. ELISA analysis of mouse urine showed a significantly higher level of CD59 (72.74±13.70 pg ml^-1^), renin (81.05±11.24 pg ml^-1^), and UMOD (1188±199.20 pg ml^-1^) in the engrafted mice compared to control mice with acellular scaffolds (Fig. 7d and Supplementary Fig. S7a,b). These proteins have been reported in adult urine and in amniotic fluid secreted by the developing kidney^32–35^. Thus, the presence of human CD59 and higher level of renin and UMOD in mouse urine indicates that the engrafted nephrons have function and produce proteins that are excreted via the lower urinary system of the recipient. These results suggest that the implanted kidney graft becomes vascularized, leads to a higher level of maturation, and achieves a degree of synthetic function after engraftment.

## Discussion

The bioengineered kidney tissues generated from hPSCs recapitulate nephron-like structures and have shown a remarkable potential to model human kidney disease^36,37^. However, several biological issues restrict their application in translational medicine. Although engrafted kidney organoids become vascularized, exhibit maturation, and have partial nephron function, they lack kidney-like architectural features such as coordinated nephron patterning, proper integration of nephron and collecting ducts, and urinary outflow^38,39^. Addressing these gaps remains critical for the advancement of kidney tissue engineering. Previous studies combining NPs and UBPs generated separately from hPSCs to form organoids ^40,41^ led us to recellularize rat native kidney scaffolds. The resulting recellularized scaffolds exhibited functional nephrons, collecting ducts and vascularization.

Native kidney scaffolds retain indispensable extracellular matrix architecture that provides a biologically active niche for recellularization, but no study showed the recellularization of kidney scaffolds using hPSC-derived NPS and UBPs^19,42^. We first evaluated if mixing both progenitors can enhance nephrogenesis and found tightly filled tubular clusters in the kidney organoids with significant upregulation of nephron markers NPHS1, LTL, CDH1, KRT8 and PECAM1. Our data is consistent with previous findings and indicate upregulation of nephron and collecting duct markers in the kidney organoids generated from mixing these progenitors^11,43,44^. The recellularization of kidney scaffolds with these mixed progenitors resulted in generation of nephron, collecting ducts, and vascularization with nephron secretory functions. Perfusion with the human endothelial marker UEA1 was detected within PECAM1 positive vascular lumens, indicating the formation of lumenized endothelial networks. Next, the detection of renin, UMOD and Vit D3 proteins in scaffold perfusion media further supports functional nephron maturation.

Analysis of *in vivo* engrafted constructs demonstrated vascularization and RBCs were detected within anastomosed human and mouse vessels. Both human and mouse vessels invaded glomerular structures in the graft and formed capillary tuft-like structures, an advanced feature of the glomerulus. Vascularization of human grafts have been reported before^45,46^ but clear anastomosis of human and mouse vessels and detection of RBCs have not been shown previously. The establishment of a functional vasculature accentuates the essential contribution of angiocrine cues and metabolic support that drive endothelial maturation *in vivo*^47^. Epithelial tubules with glomerulus, proximal tubules, loop of Henle, distal tubules, and collecting ducts were detected in the human graft. Further staining with HuNu suggests that nephron, collecting ducts and stromal cells in the graft have a human origin. PDGFRβ expressing stromal cells were detected between epithelial tubules, similar to previous reports^26,48^. Additionally, loss of transcription factor PAX2 induced nephron progenitors adapt stromal cell-like identity in the developing kidney^49^, suggesting this expansion is associated with graft micro-environmental cues. To maintain secretory function and to ensure proper filtrate excretion, the mammalian urinary system depends on the integration of nephrons with the renal collecting duct network and subsequent fusion of the ureteric extension with the bladder, regulated by Ret and retinoic acid signaling^50,51^. Engrafted recellularized scaffolds generated nephron segments connected in sequence, including glomeruli linked to LTL positive proximal tubules, CDH1 expressing loops of Henle, UMOD positive thick ascending limbs, NCC positive distal tubules, and DBA positive collecting ducts, suggesting that these nephrons contain interconnected segments with collecting ducts within the graft. Furthermore, the detection of tubule and human specific proteins in mouse urine strengthen the evidence for interconnected nephrons with the collecting duct. Notably, nephron–collecting duct fusion was not observed *in vitro*, suggesting that *in vivo* cues are required to drive this integration.

The limitation of this study is that we were unable to visualize a complete nephron extending fully into a collecting duct due to the technical complexity of the investigation. Additionally, while human proteins were detected in mouse urine, their presence could result from diffusion or active tubular flow. However, the observed structural continuity strongly suggests the latter. In this study, the engrafted scaffolds were evaluated after three weeks of engraftment, though a long-term evaluation is required to study the fate of engrafted tissue *in vivo*.

In summary, we demonstrate that human hPSCs-derived NPs and UBPs can reconstitute acellular kidney scaffolds and lead to functional and mature renal tissue. To our knowledge, this is the first study to report a kidney scaffold reconstituted with mixed progenitors that mature within a bioreactor, leading to engraftment with functional anastomoses to host circulation. The perfusable scaffold enables patterning, maturation and function of nephron and collecting ducts with the formation of lumenized endothelial networks that fused with host vessels after engraftment, evidenced by the presence of mouse erythrocytes within host- and graft-derived lumens. These findings establish a foundational platform for evaluating the therapeutic potential of bioengineered kidney tissues in disease models, with future studies directed toward clinical applications.

## Methods

### Cell culture and maintenance

Use of the human embryonic PSCs line H9 (WiCell, WA09, NIH approval number NIHhESC-10-0062)^52^ was approved by the ESCRO committee at the University of Arizona. These cells were cultured and maintained in feeder-free conditions on geltrex (ThermoFisher) coated 6-well plate in StemFit Basic03 medium (amsbio) in a 37°C incubator with 5% CO_2_. For routine maintenance, 70%-80% confluent hPSCs colonies were passaged at a split ratio 1:10 using Gentle Cell Dissociation Reagent (STEMCELL Technologies) following manufacturer’s protocol.

### Animal experiments

Lewis rats (250–300 g) were purchased from Charles river laboratories and used to obtain native kidneys for decellularization into scaffolds. Male NOD.Cg-Prkdcscid Il2rgtm1Wjl/SzJ (NSG) mice (15-25g) were purchased from Jackson laboratory for engraftment experiments. All animal experiments were reviewed and approved by the Institutional Animal Care and Use Committees (IACUC) of Northwestern University (IS00003195) and/or the University of Arizona (#2020-0677). All *in vivo* experiments were conducted in compliance with the ARRIVE Guidelines 2.0. The manuscript has been prepared in accordance with the ARRIVE Essential 10 checklist.

### Differentiation of hPSCs into nephron and ureteric bud progenitors

To differentiate hPSCs into NPs, cells were dissociated into single cells using Accutase (STEMCELL Technologies) and plated at a cell density 1.8×10^5^ cells per well on geltrex coated 6-well plate in StemFit Basic03 medium. We followed the Morizane et al. 2015 protocol to differentiate hPSCs into NPs^23^. In brief, once cell cultures reached ∼50% confluence, cells were treated with 8μM CHIR99021 (Reprocell) for 4 days, 10ng/ml ActivinA (R&D Systems) for three days, and 10ng/ml FGF9 (R&D Systems) in basic medium Advanced RPMI1640 and 1×GlutaMAX (ThermoFisher) for the next two days. After 9 days of differentiation, nephron progenitors were dissociated with TrypLE Express (ThermoFisher), characterized, and used for organoid generation or seeding in acellular kidney scaffolds.

We followed the Tsujimoto et al. 2020 published protocol to differentiate hPSCs into UBPs with slight modifications^24^. In brief, cells were dissociated into single cells using Accutase (STEMCELL Technologies) and plated at a cell density 2.5×10^5^ cells per well on geltrex coated 6-well plate in StemFit Basic03 medium. After 24h of culture, cells were treated with 100 ng/ml activin A, 10 ng/ml BMP4 (R&D Systems) and 3μM CHIR99021 in a basal medium CTS Essential 6 Medium (ThermoFisher). After 24h of treatment, differentiating cells were treated with 200 ng/ml FGF8 (R&D Systems), 0.1μM 4-[(E)-2-(5,6,7,8-Tetrahydro-5,5,8,8-tetramethyl-2-naphthalenyl)-1-propenyl] benzoic acid (TTNPB, R&D Systems), 1μM A83-01 (Millipore Sigma) and 0.1μM LDN193189 (Reprocell) for 3 days. Next, cells were treated with 200 ng/ml FGF8, 0.1μM TTNPB, 0.1 µM LDN, 100 ng/ml glial cell line-derived neurotrophic factor (GDNF, R&D Systems) and 1µM CHIR99021 for 2 days. After 6 days of differentiation, UB progenitors were dissociated with TrypLE Express, characterized and used for organoid generation or seeding in acellular kidney scaffolds.

### Cryopreservation and revival of progenitors

To cryopreserve NPs and UBPs, cells were harvested with TrypLE Express and single cells were resuspended 4-5 million cells ml^-1^ ice-cold NutriFreez D10 Cryopreservation Medium (Biological Industries). Then Nunc cryogenic tubes (ThermoFisher) containing 1 ml cell suspension in each vial was transferred to a Corning CoolCell box (Corning) and frozen at -80°C overnight. After 24h, frozen vials were transferred to liquid nitrogen. To revive cryopreserved progenitors, vials were swirled gently in 37°C water bath and resuspended in 5 ml Advanced RPMI1640 and 1×GlutaMAX for NPs or CTS Essential 6 Medium for UBPs containing 10% knockout serum supplement (ThermoFisher) and 10 µM Y-27632 (Millipore Sigma). The cell suspension was centrifuged and cell supernatant was discarded. The thawed NPs were plated on geltrex coated 6-well plate in Advanced RPMI1640 containing 1×GlutaMAX, 10ng/ml FGF9 and 10 µM Y-27632. Thawed UBPs were plated on geltrex coated 6-well plate in CTS Essential 6 Medium containing 200 ng/ml FGF8, 0.1μM TTNPB, 0.1 µM LDN, 100 ng/ml GDNF, 1µM CHIR99021 and 10 µM Y-27632. Cells were harvested after 24h of culture and used in differentiation. We successfully revived and tested NPs and UBPs that had been cryopreserved for more than 12 months without any deprivation of differentiation potential.

### Generation of kidney organoids by combining NPs and UBPs

Before combining NPs and UBPs, differentiation of hPSCs into NPs began three days prior (day -3) to the differentiation of hPSCs into UBPs; the initiation of UBP differentiation is referenced at day 0 and this time scale is used to provide a synchronous reference frame to describe NP and UBP differentiation. On day 6, NPs (at their 9^th^ day of differentiation) and UBPs (at their 6^th^ day of differentiation) were combined together at a 3:1 ratio^53^ and aggregated at 5.0×10^5^ cells/aggregate at the air-liquid interface^8^ in organoid media containing 100ng/ml FGF9, 100ng/ml BMP7 (R&D Systems), 1μg/ml Heparin (Millipore Sigma) and 0.1μM TTNPB in a basic medium APEL2 (Stem cell Technologies). To aggregate cells, an air-liquid interface was created by floating Isopore Membrane Filter (Millipore Sigma) at 1 ml organoid media in a 24-well plate. To generate an NP-only lineage control to compare the effect of mixed progenitors on nephrogenesis within kidney organoids, only NPs were aggregated at 5.0×10^5^ cells/aggregate at the air-liquid interface in the same organoid media composition. Organoid media was replaced with fresh media every 48 h of culture. On day 6 of aggregation, 50 ng/ml EGF (R&D Systems) was supplemented in the media to enhance epithelialization^54–56^. On day 10 of aggregation, kidney organoids were harvested for evaluation.

### Immunofluorescence staining

NPs and UBPs were fixed with 4% paraformaldehyde at room temperature (RT) for 7 min. Cells were permeabilized with 0.3% Triton X-100 for 10 min at RT and incubated 1h at RT with blocking buffer consisting of 5% donkey serum in PBS. Primary antibodies listed in Supplementary Table 1 were diluted in blocking buffer and cells were incubated with diluted primary antibodies overnight at 4°C. The next day, cells were washed two times with PBS and incubated at RT for 1h with secondary antibodies diluted in blocking buffer and containing 0.01 µg ml^-1^ 4′,6-diamidino-2-phenylindole (DAPI). Next, cells were washed two times with PBS and imaged at Axio Observer microscope (Zeiss Microscopy).

### RNA isolation and real time PCR

Total RNA was isolated from differentiating cells or kidney organoids using PureLink RNA Mini Kit following the manufacture’s instruction. Isolated RNA was quantified on the Nanodrop One Spectrophotometer (Thermo Fisher Scientific) and 2 µg of total RNA was reverse transcribed with a High-Capacity cDNA Reverse Transcription Kit (Applied Biosystems) according to the manufacturer’s instructions. Quantitative polymerase chain reaction was performed with SsoAdvanced Universal SYBR Green Supermix (Bio-Rad) on a CFX connect real-time PCR system (Bio-Rad). Relative fold expression levels were analyzed by the ΔΔCT method and normalized to TATA-Box Binding Protein (TBP) gene expression. All primers used in Real Time PCR experiments are listed in Supplementary Table 2.

### Native rat kidney recovery and decellularization

Rat kidneys were recovered and decellularized following our previously published protocol^27^. Male or female Lewis rats were anesthetized using intraperitoneal injection of 50 mg/kg pentobarbital (MWI Animal Health), and a longitudinal midline incision was performed to open the abdomen. The abdominal aorta and the inferior vena cava were dissected from above the right renal hilum to the iliac bifurcation, and the ureter was dissected from the urinary bladder. Then after 2,000 USP heparin units/kg of body weight (Millipore Sigma) was injected through the dorsal penile vein, the abdominal aorta was ligated above the right renal artery and cannulation of the infrarenal abdominal aorta was done using an 18G cannula (BD biosciences). Both rat kidneys were gently perfused with 10 ml cold saline through the infrarenal aortic cannula. The right renal artery and vein were transected and the right kidney was removed after perfusion. Then the left kidney along with the attached aortic cannula was removed after the suprarenal abdominal aorta and the left renal vein were transected. Kidneys were decellularized by sequential perfusion of solution (Fig. 2b and Supplementary Fig. S3). In brief, kidneys were first connected with one end of a silicone rubber tubing and suspended in a perfusate reservoir and other end was immersed in a reagent reservoir. A peristaltic pump (Cole-Parmer) was connected at the middle of the tubing. Each kidney was perfused with 500 ml distill water (dH_2_O) at 5 ml/min for 1h, 40 min, 1000 ml 1% Triton X-100 (Millipore Sigma) at 5 ml/min for 3h, 20 min, 1000 ml 1% Triton X-100 at 1 ml/min for 16h, 40 min, 1000 ml 0.1% SDS (Millipore Sigma) at 5 ml/min for 3 hr, 20 min. Finally, decellularized kidneys were washed with 500 ml dH_2_O at 5 ml/min for the next 1h, 40 min. Decellularized kidneys were validated for the effectiveness of decellularization by quantification of extracted residual DNA from kidney scaffolds as a surrogate marker for cell removal. DNA from scaffolds and native kidney was extracted using DNeasy Blood & Tissue Kit (Qiagen) following manufacturer’s instruction. Extracted DNA was quantified on the Nanodrop One Spectrophotometer. The extracted residual DNA from kidney scaffolds (n=6) was compared with extracted DNA from native rat kidneys (n=6) and data represented as DNA content (ng/mg of wet tissue).

### Recellularization of acellular kidney scaffolds

To recellularize acellular kidney scaffolds with NP and UBP lineage cells, the kidney scaffolds were placed in a glass bioreactor (Fig. 2d and Supplementary Fig. S4) that we have described^6^ and perfused with 50 ml sterilization solution containing 0.1% peracetic acid (Millipore Sigma), 4% ethanol (Millipore Sigma) in autoclaved dH2O for 30 min. After sterilization, the kidney scaffolds were washed with PBS (Thermo Fisher Scientific) three times for 30 min each. Then, kidney scaffolds were perfused with 40 ml recellularization media containing 100ng/ml FGF9, 100ng/ml BMP7, 1μg/ml Heparin and 0.1μM TTNPB in a basic medium APEL2 for 1h. Next, NPs and UBPs were mixed together at a ratio 3:1 and seeded through the cannulated renal artery at a cell density 100 million cells in each kidney scaffold. Immediately after cell seeding, the flow rate was increased to 25ml min^-1^ for the next 15 min for a homogenous distribution of cells and stopped, similar to our described method.^7^ The bioreactor assembly was then transferred to a 37°C incubator with 5% CO_2_ incubator and left undisturbed for at least 6h to allow attachment of seeded cells. Thereafter, flow was resumed with a perfusion flow rate of 1ml min^-1^ for the next 10 days to recellularize the scaffold. The recellularization media was replaced with fresh media every 48 h of culture. On day 6 of recellularization, 50 ng/ml EGF (R&D Systems) was supplemented in the media to enhance epithelialization. On day 16 of differentiation, recellularized kidney scaffolds were removed from the bioreactor for evaluation or engraftment. To evaluate cell viability in kidney scaffolds, 3-(4,5-dimethylthiazol-2-yl)-2,5-diphenyltetrazolium bromide (MTT) assay was performed using commercial kit (Abcam). Mixed progenitors (NPs and UBPs, 3:1) were seeded with recellularization media in a culture plate without a scaffold (No scaffold, n=6), in the kidney scaffold (with scaffold, n=6) and in the kidney scaffold with 1% sodium dodecyl sulfate (SDS) as a negative control (n=6). 24h after seeding, MTT assay was performed following manufacturer’s instruction and absorbance was taken at 590 nm using a Microplate Reader (Biotek).

Hematoxylin and eosin staining was performed on recellularized kidney scaffolds to evaluate degree of recellularization. To evaluate AQP2 translocation, the perfusion media on day 14 was supplemented with 10 nM AVP and 10 nM Aldo for 48h. The recellularized scaffolds were fixed and stained for AQP2. Next, recellularized scaffolds were also evaluated for the presence lumenized endothelial networks. We perfused recellularized scaffold with 5 µg/ml DyLight594-conjugated UEA1 (Vector Laboratories) in recellularization media for 1 hour at a flow rate 1ml min^-1^ in a 37°C incubator. Following perfusion, scaffolds were washed three times with PBS and stained for endothelial markers. Fluorescent images were acquired at Zeiss LSM880 inverted confocal microscopes (Zeiss Microscopy).

### *In vivo* engraftment of recellularized kidney scaffolds

Male NSG mice were divided in two groups; acellular scaffold engraftment group (control, n=6) and recellularized scaffold engraftment group (engraftment, n=6). Healthy NSG mice served as healthy control (healthy, n=6). Mice underwent 5% inhaled isoflurane (MWI Animal Health) induction and were maintained with ventilated 1–3% inhaled isoflurane throughout the survival surgery. The mouse was placed on a heating pad on its right recumbent position, and an incision was made in the left flank. The left kidney was externalized, and the kidney pedicel was clamped with an atraumatic clamp. The mouse adrenal gland was gently pushed aside with forceps to expose the kidney pole. Then, ∼3mm tissue from upper pole of the left kidney was removed and ∼3mm of recellularized kidney scaffold, after washing with PBS, was sutured to the cut surface of the kidney using 6-0 vicryl suture (Ethicon). The cellularized scaffold was wrapped with a small piece of surgifoam (Ethicon) and the vascular clamp was removed with gentle pressure on surgifoam to keep it intact. Once bleeding was controlled after reperfusion, the surgifoam was removed gently and kidney was returned to the abdominal cavity. The body cavity was closed with 4-0 vicryl suture and the skin was closed using 9mm autoclips (Mikron Precision). Three weeks after engraftment, mouse kidneys were recovered for further evaluation.

### Whole mount immunofluorescence staining

To stain kidney organoids and 100µm vibratome engrafted kidney sections, whole mount immunofluorescence staining was performed. Kidney organoids were fixed with 4% paraformaldehyde for 15 min at RT and then permeabilized with 1% Triton X-100 in PBS for 10 min at 4 °C. Engrafted kidneys were pared down so that only a transverse slab of the kidney underlying the graft remained and this tissue was fixed with 4% paraformaldehyde for 1h mm^-1^ of tissue thickness at 4°C before vibratome sectioning at 100 µm. Vibratome sections were permeabilized with 1% Triton X-100 in PBS for 10 min at 4°C. Both kidney organoids and vibratome sections were blocked and diluted in blocking buffer containing 5% donkey serum in PBS for 1h at 4°C. Next, blocked tissues were incubated with primary antibodies listed in Supplementary Table 1 diluted in blocking buffer at 4°C overnight. The following day they were washed with PBS for 6h at 4°C and incubated with secondary antibodies containing 0.01 µg ml^-1^ DAPI at 4°C overnight. The next day after washing with PBS for 6h at 4°C, stained kidney organoids or vibratome tissue sections were mounted in Vectashield antifade mounting medium (Vector Laboratories) and imaged at Zeiss LSM880 inverted confocal microscopes (Zeiss Microscopy). Percent area fraction (a.f.) for different markers in kidney organoids was measured using Image J. To quantify area fraction for each antibody stain, five separate fields from six different specimens were measured and the mean was plotted with standard deviation (mean ± s.d.).

### Protein evaluation by Enzyme Linked Immunosorbent Assay and colorimetric assays

The decellularized rat kidney scaffolds (n=6) were evaluated for the ECM components and compared with native rat kidneys (n=6). Total Glycosaminoglycans assay kit (Abcam) and rat fibronectin Enzyme Linked Immunosorbent Assay (ELISA) kit (Novus Biologicals) were used. For both the assays, 25 mg tissues were processed following manufacturer’s instructions and assays were performed following manufacturer’s protocol. Next, secreted proteins in recellularized scaffold perfusion media were analyzed by ELISA. To evaluate conversion of 25-hydroxyvitamin D3 into 1, 25-dihydroxyvitamin D3, recellularized scaffold on day 16 was perfused with recellularization media containing 100 ng ml^-1^ 25-hydroxyvitamin D3 (Millipore Sigma). After 24h of perfusion, the recellularized scaffold media was collected and stored at -80°C. Secreted proteins Renin (R&D systems), UMOD (R&D systems) and 1, 25-dihydroxyvitamin D3 (My BioSource) in perfusion media were evaluated by ELISA using the manufacturer’s instructions. To evaluate secreted proteins in mouse urine, ELISA was performed. Mice urine was recovered from all the experimental groups on day 7 and day 21 after implantation and centrifuged at 5000× g for 5 min. Urine supernatant was aliquoted and stored in -80°C freezer until use. Secreted proteins Renin, UMOD and CD59 (Abcam) in mouse urine were evaluated by ELISA kits using the manufacturer’s instructions. Urine from healthy mice and mice with engrafted acellular scaffolds served as healthy and control urine respectively.

### Statistics

All experiments were performed at least three times, with one representative dataset shown. Data are presented as mean ± s.d. GraphPad Prism version 8.0 (GraphPad Software) was used for statistical analysis. Statistical analyses between two different groups were performed by using Student t-test and one-way ANOVA followed by Tukey’s multiple-comparison test was used for multiple group comparison. Differences with values of *P*<0.05 were considered statistically significant. Sample sizes are provided in the figure legends.

### Data availability

Data supporting the study are available within the paper and its supplementary information. Any additional data or materials will be available upon request to the corresponding author.

## Acknowledgements

Surgical recovery of rat kidneys was performed by the Microsurgery and Preclinical Research Core at Northwestern University Comprehensive Transplant Center. The authors would like to acknowledge and thank the (Re)Building a Kidney Consortium of the National Institute of Diabetes and Digestive and Kidney Diseases of the National Institutes of Health in addition to Patty Jansma and Doug Cromey for assistance with fluorescent imaging at the University of Arizona Imaging Cores - Optical Facility.

## Funding information

J.A.W. discloses support for this work from the University of Arizona to include *in vivo* studies, the National Institute of Diabetes and Digestive and Kidney Disease (NIDDK) of the National Institutes of Health [grant number R01DK113168] and Merit Review Award from the United States Department of Veterans Affairs Biomedical Laboratory Research and Development Service [grant number I01BX002660]. Z.J.Z. discloses support for this work from Robert R. McCormick Foundation. The content is solely the responsibility of the authors and do not represent the views of the U.S. Department of Veterans Affairs, the National Institutes of Health or the United States Government.

## Author contributions

A.K.G. and J.A.W. developed the study and experimental design. A.K.G. and E.M. performed and analyzed experiments. J.J.W. and Z.T. recovered rat kidney for decellularization work. All authors wrote and/or edited the manuscript. J.A.W. and Z.J.Z. acquired funding for this work. All authors have reviewed and approved the final manuscript.

## Competing interests

The authors declare no competing interests.

## Notes

### Competing Interest Statement

The authors have declared no competing interest.

