## Supplementary Materials for "Coalescing nephron and ureteric bud progenitors potentiates nephrogenesis in recellularized kidney scaffolds"

**Supplementary data:**

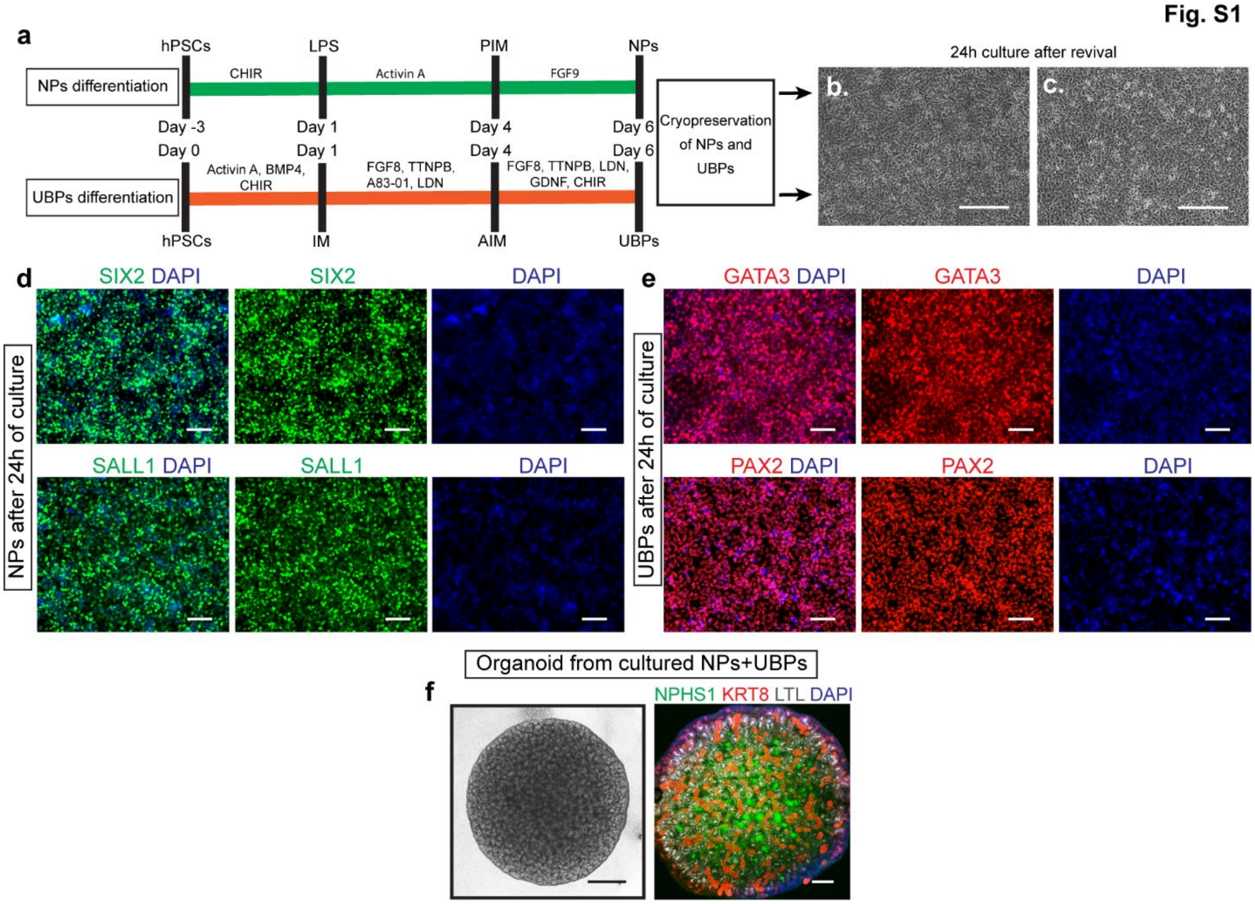


**Fig. S1 | Cryopreserved NPs and UBPs express progenitor markers and retain nephrogenic potential. a.** Schematic diagram shows the stepwise differentiation and cryopreservation of NPs and UBPs. **b.** Revived NPs and **c.** UBPs after 24h of culture. Scale bars 100µm. **d.** Representative immunofluorescence images show expression of progenitor markers SIX2 and SALL1 by NPs. Scale bars 100µm. **e.** Representative immunofluorescence images shows expression of progenitor markers GATA3 and PAX2 by UBPs. Scale bars 100µm. **f.** Representative images show generation of kidney organoids (scale bar 500µm) from a combination of cryopreserved NPS and UBPs cells and immunofluorescence staining (scale bars 100µm) reveals expression of NPHS1 expressing podocytes, KRT8 expressing epithelial tubules and LTL positive proximal tubules. Scale bars 100µm.

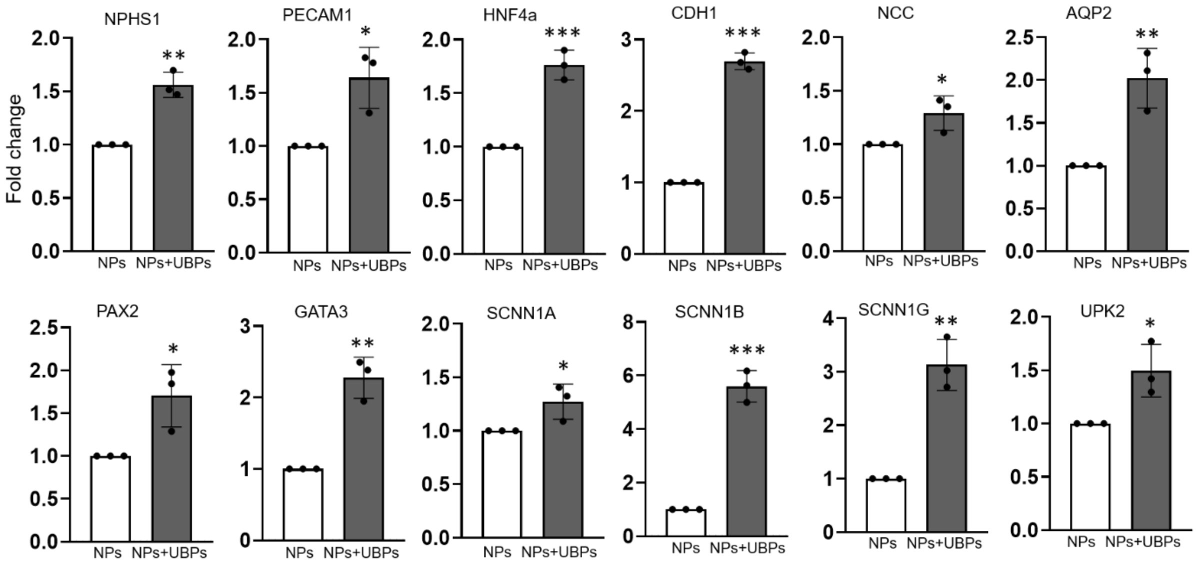


**Fig. S2 | Gene expression analysis by qPCR in kidney organoids generated from NPs only or by a combination of NPs and UBPs.** Kidney organoids generated by combining NPs and UBPs have significant upregulation of genes for podocytes (NPHS1), endothelial networks (PECAM1), proximal tubules (HNF4α), epithelial tubules (CDH1), distal tubules (NCC), collecting ducts (PAX2 and GATA3), collecting duct principle cells (AQP2, SCNN1A, SCNN1B and SCNN1G), and urothelial gene (UPKII) in comparison to kidney organoids generated from NPs only ($n = 6$ organoids from three independent biological replicates per group). *$P < 0.05$; **$P < 0.01$; ***$P < 0.001$.

Fig. S3

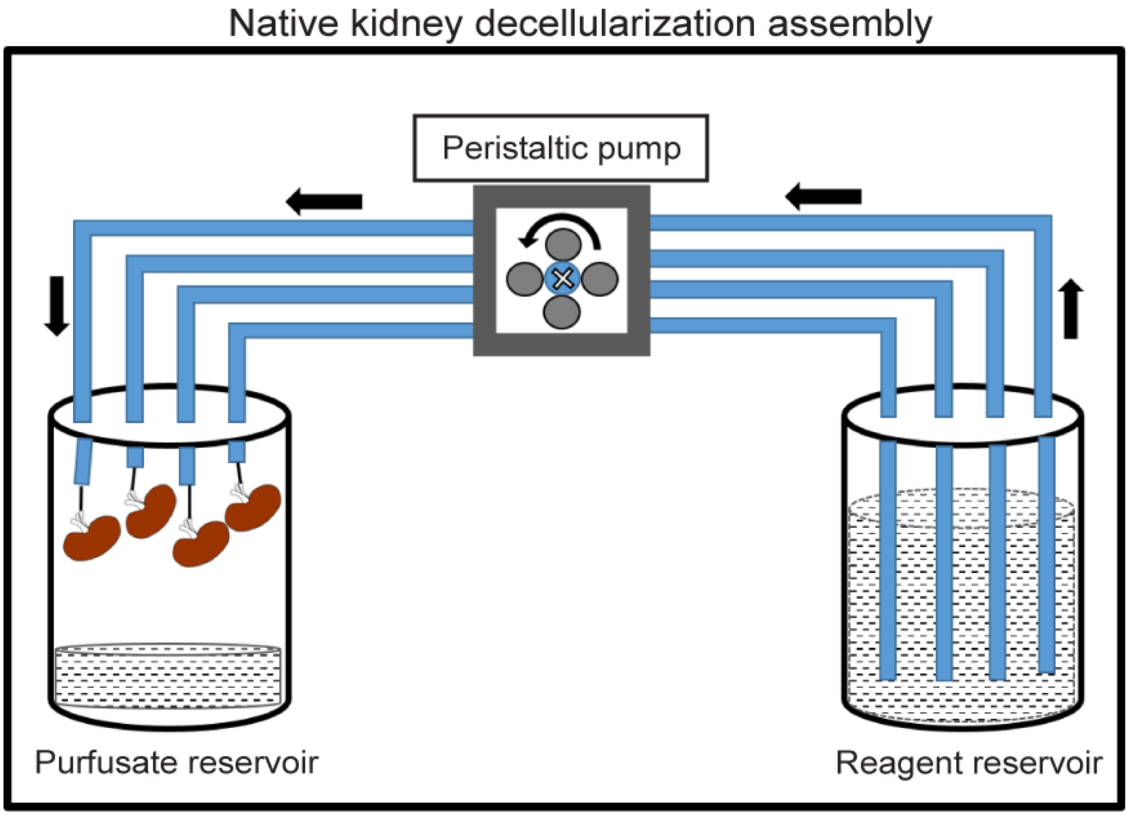


**Fig. S3 | Decellularization of rat native kidneys.** Schematic diagram depicts the decellularization assembly to develop kidney scaffolds.

Fig. S4

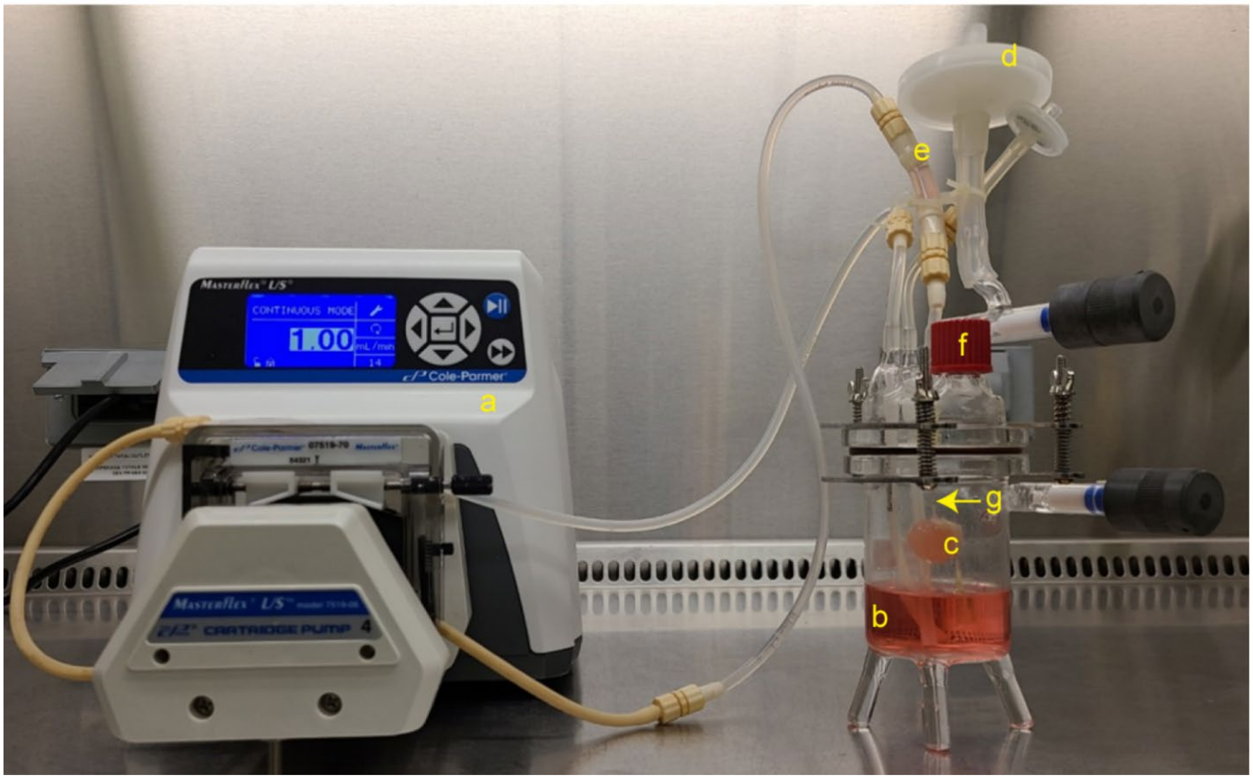


**Fig. S4 | Recellularization assembly.** During recellularization **a.** peristaltic pump was connected to **b.** glass bioreactor and the **c.** kidney scaffold was perfused with media in a bioreactor. **d.** 0.2µm filter **e.** Air-bubble trapper **f.** Outlet for media change **g.** Renal artery.

Fig. S5

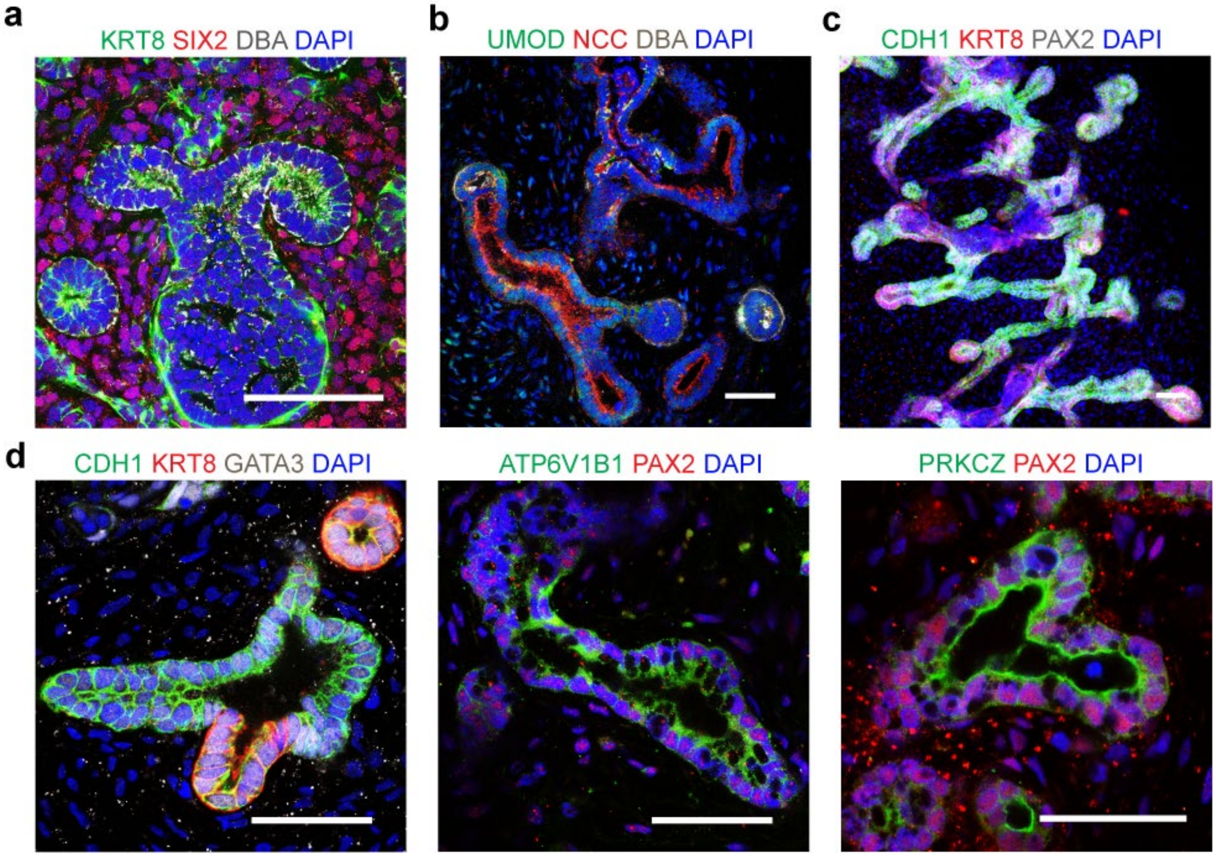


**Fig. S5 | Combining NPs and UBPs resulted in formation of nephrogenic zone-like structures and advanced collecting ducts. a.** Representative immunofluorescence images show SIX2$^+$ nephron progenitors around KRT8$^+$ and DBA$^+$ ureteric buds or collecting ducts in the recellularized scaffold. These images indicate that nephrogenic zone-like structure still exist even after ten days of recellularization. **b.** Tubules showing segmentation in UMOD$^+$ loop of Henle, NCC$^+$ distal tubule and DBA$^+$ collecting ducts in the graft. **c.** Arborized collecting duct-like structures in the graft. **d.** Collecting ducts also expressed CDH1, KRT8, GATA3, PRKCZ, PAX2 and showed presence of ATP6V1B1 expressing intercalated cells. Scale bars 50µm.

Fig. S6

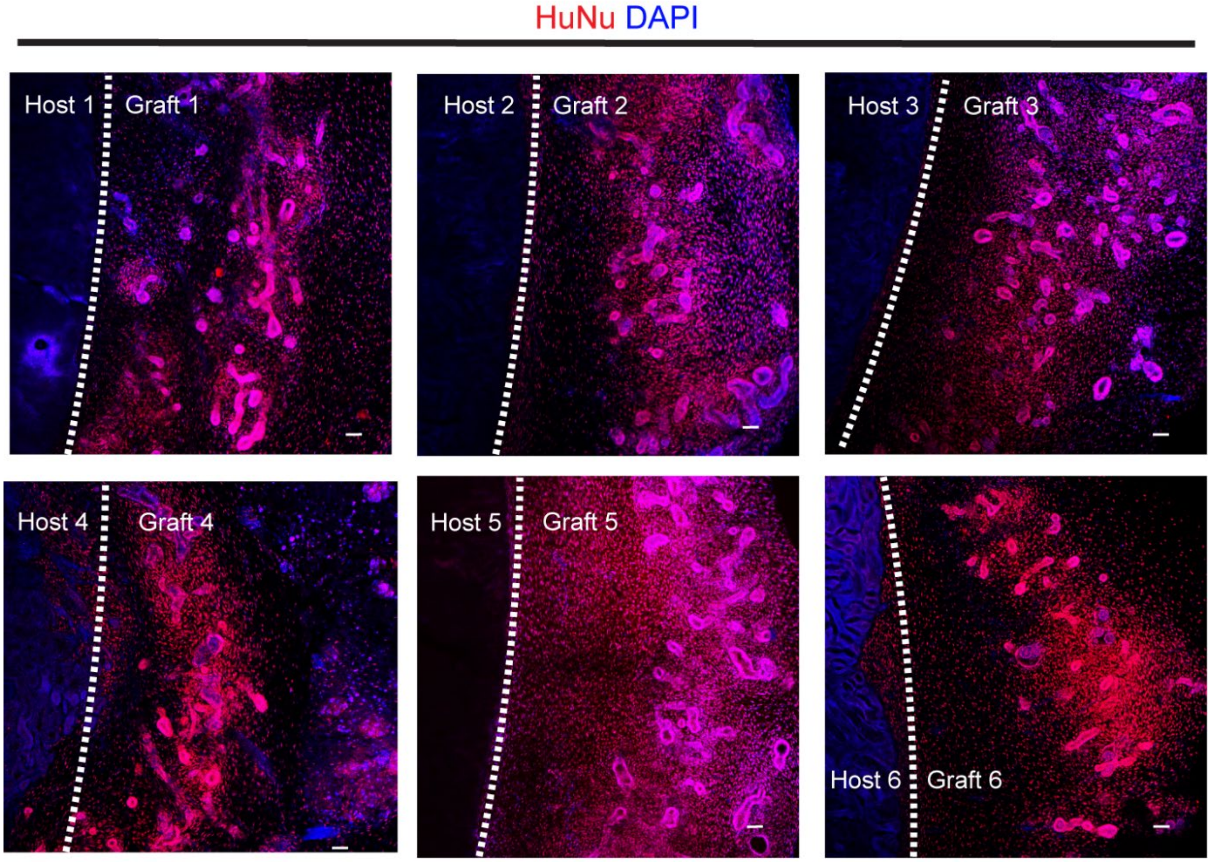


**Fig. S6 | Graft characterization for human nuclear antigen.** Tissue sections of host and graft tissue were stained with human nuclear antigen (HuNu) to determine the origin of cells within the graft. Representative immunofluorescence images of tissues sections from all six mice show expression of HuNu whereas mouse kidney tissues do not have any expression of HuNu. HuNu expression indicates the graft survived, including areas remote from the interface with the cut surface of the host kidney. Scale bars 50µm.

**Fig. S7**

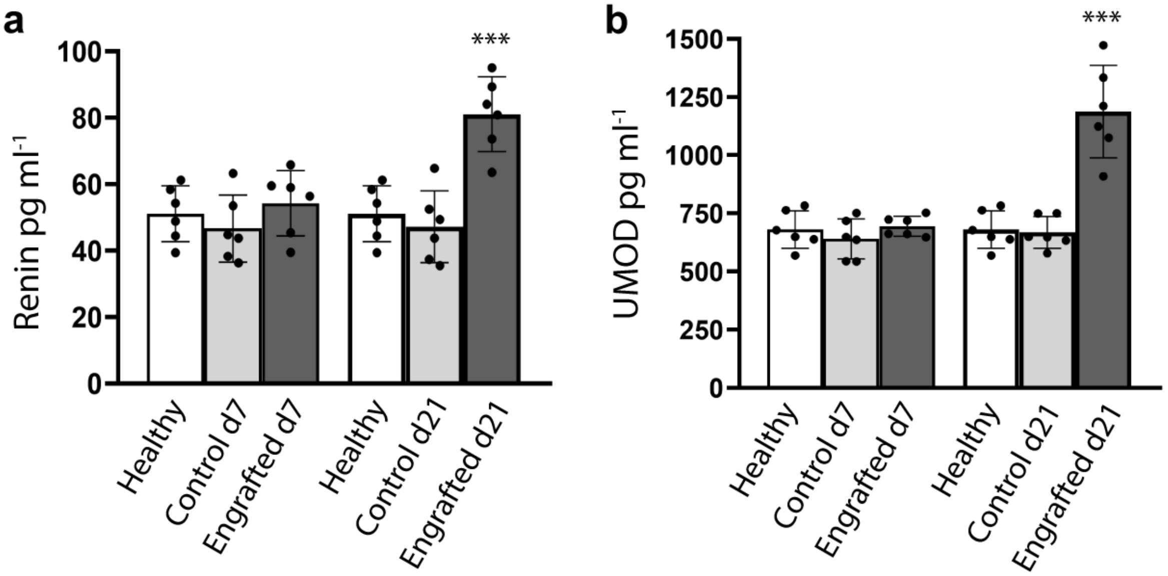


**Fig. S7 | Detection of renin and uromodulin in mouse urine by ELISA.** Proteins **a.** renin and **b.** uromodulin (UMOD) were detected at a significantly higher level using a human-specific antibody in the urine of engrafted mice on day (d) 21 after implantation in comparison to mice implanted with an acellular scaffold (Control) or healthy mice, (n = 6 mouse per group from three independent biological replicates). Renin and uromodulin proteins were also detected in mouse urine using this human-directed antibody. These proteins share higher percentage of sequence similarity between mouse and human, suggesting that the significant increase in the d21 engrafted group is likely to be due to the engrafted human tissue. Data presented as mean ± s.d. ***$P < 0.001$.

**Supplementary Table 1 |** List of antibodies/lectins.

| **Name** | **Species** | **Manufacture** | **Catalog number** | **Dilution** |
|---|---|---|---|---|
| OSR1 | Rabbit | Cell signaling Tech | 3729S | 1:100 |
| HOXD11 | Mouse | MilliporeSigma | SAB1403944 | 1:100 |
| LHX1 | Mouse | DSHB | 4F2-c | 1:100 |
| SIX2 | Rabbit | Proteintech | 11562-1-AP | 1:100 |
| SALL1 | Mouse | Novus Biologicals | PP-K9814-00 | 1:100 |
| WT1 | Rabbit | Abcam | ab89901 | 1:100 |
| GATA3 | Rabbit | Abcam | ab199428 | 1:100 |
| PAX2 | Rabbit | Biolegend | 901001 | 1:100 |
| PRKC ζ | Mouse | Santa Cruz | sc17781 | 1:100 |
| ETV5 | Rabbit | Abcam | ab102010 | 1:100 |
| NPHS1 | Sheep | R&D systems | AF4269 | 1:100 |
| CDH1 | Goat | R&D systems | AF648 | 1:100 |
| CDH1 | Rabbit | Abcam | ab11512 | 1:100 |
| LTL | - | Vector Lab. | B-1325-2 | 1:200 |
| KRT8 | Rat | MilliporeSigma | MABT329 | 1:100 |
| PECAM1 | Mouse | Cell signaling Tech | 3528S | 1:50 |
| PECAM1 | Rat | BD Pharmingen | 550274 | 1:50 |
| POU3F3 | Rabbit | Invitrogen | PA5-64311 | 1:100 |
| CALB1 | Mouse | MilliporeSigma | C9848 | 1:100 |
| HOXB7 | Rabbit | Cell signaling Tech | 65130S | 1:100 |
| RET | Rabbit | Thermo Scientific | MA5-32342 | 1:100 |
| DBA | - | Vector Lab. | B-1035-5 | 1:200 |
| PODXL | Goat | R&D systems | AF1658 | 1:100 |
| RENIN | Rabbit | Abcam | ab212197 | 1:100 |
| NaKATPase | Rabbit | Abcam | ab76020 | 1:100 |
| UMOD | Sheep | MilliporeSigma | AB733 | 1:100 |
| UPKII | Rabbit | Novus Biologicals | NBP2-33389 | 1:100 |
| UEA1 | - | Vector Lab. | DL-1067-1 | 1:200 |

| | | | | |
|---|---|---|---|---|
| AQP2 | Rabbit | MilliporeSigma | 178612 | 1:100 |
| ATP6V1B1 | Mouse | Santa Cruz | Sc55544 | 1:100 |
| RBC | Rabbit | Rockland | 110-4139 | 1:100 |
| PDGFRβ | Rabbit | Abcam | ab32570 | 1:100 |
| LRP2 | Mouse | MilliporeSigma | MABS489 | 1:100 |
| NCC | Rabbit | MilliporeSigma | HPA028748 | 1:100 |
| NPHS2 | Rabbit | Novus Biologicals | NBP2-75624 | 1:100 |
| NKCC2 | Rabbit | Abcam | ab171747 | 1:100 |
| Ac-αTubulin | Mouse | Proteintech | 66200-1-Ig | 1:100 |
| SCNN1A | Rabbit | Alomone | ASC-030 | 1:100 |
| SCNN1B | Rabbit | Proteintech | 14134-1-AP | 1:100 |
| SCNN1G | Rabbit | Proteintech | 13943-1-AP | 1:100 |
| HuNu | Mouse | MilliporeSigma | MAB1281B | 1:50 |

**Supplementary Table 2 |** List of qPCR primer sequences.

| Gene | Forward primer | Reverse primer |
|---|---|---|
| AQP2 | CACGTCTCCGTTCTCCGAG | CTGTTGCTGAGAGCATTGACA |
| CHD1 | CACCCTGGCTTTGACGCCGA | AAACGGAGGCCTGATGGGGCG |
| GATA3 | GCCCCTCATTAAGCCCAAG | TTGTGGTGGTCTGACAGTTCG |
| GDNF | GGCAGTGCTTCCTAGAAGAGA | AAGACACAACCCCGGTTTTTG |
| HNF4α | CGAAGGTCAAGCTATGAGGACA | ATCTGCGATGCTGGCAATCT |
| NANOG | TGATTTGTGGGCCTGAAGAAA | GAGGCATCTCAGCAGAAGACA |
| NCC | TGGACGACCATTTCCTACCTGG | CACTCGGTGAAGTTCCAGCCAT |
| NPHS1 | CTGCCTGAAAACCTGACGGT | GACCTGGCACTCATACTCCG |
| OCT4 | CCTGAAGCAGAAGAGGATCACC | AAAGCGGCAGATGGTCGTTTGG |
| PAX2 | TGTCAGCAAAATCCTGGGCAG | GTCGGGTTCTGTCGTTTGTATT |
| PECAM1 | AACAGTGTTGACATGAAGAGCC | TGTAAAACAGCACGTCATCCTT |
| SCNN1A | GTGCCTACATCTTCTATCCGCG | GTCTGAGGAGAAGTCAACCTGG |
| SCNN1B | AGACAACCACAATGGCTTAACA | TGAGGCTACATAGTCTCATGGC |
| SCNN1G | GCACCCGGAGAGAAGATCAAA | TACCACCGCATCAGCTCTTTA |
| SIX2 | AAGGCACACTACATCGAGGC | CACGCTGCGACTCTTTTCC |
| T | CCTTCAGCAAAGTCAAGCTCACC | TGAACTGGGTCTCAGGGAAGCA |
| TBP | TGTATCCACAGTGAATCTTGGTTG | GGTTCGTGGCTCTCTTATCCTC |
| TBX6 | TCATCTCCGTGACAGCCTACCA | CCGCAGTTTCCTCTTCACACGG |
| UPK2 | CACTGAGTCCAGCAGAGAGATC | ACAGAGAGCAGCACCGTGATGA |
| WNT9B | TGTGCGGTGACAACCTCAAG | ACAGGAGCCTGATACGCCAT |

**Supplementary Table 3 |** Genes/markers to identify nephron segments/function.

| Gene Name | Identification marker/Function |
|---|---|
| OCT4, NANOG | Pluripotency marker |
| T, TBX6 | Mesoderm marker |
| OSR1 and HOXD11 | Posterior intermediate mesoderm |
| OSR1 and LHX1 | Anterior intermediate mesoderm |
| SIX2, GDNF, SALL1, WT1 | Nephron progenitors |
| GATA3, WNT9B, PAX2, ETV5 | Ureteric bud progenitors |
| NPHS1, WT1, PODXL, NPHS2 | Podocytes |
| CDH1 | Epithelial cells |
| PECAM1, UEA1 | Endothelial cells |
| CALB1, PAX2, HOXB7, RET, PRKCζ, KRT8 | Collecting ducts |
| AQP2, SCNN1A, SCNN1B, SCNN1G, | Principal cells |
| ATP6V1B1 | Intercalated cells |
| LTL, NaKATPase, LRP2, HNF4α | Proximal tubule |
| NKCC2, UMOD | Loop of Henle |
| POU3F3, NCC | Distal tubule |
| Ac-αTubulin | Microtubules |
| UPKII | Urothelial cells |
| PDGFRβ | Stromal cells |
| Renin | Juxtaglomerular cells |
